# Characterization of cell-to-cell variation in nuclear transport rates and identification of its sources

**DOI:** 10.1101/001768

**Authors:** Lucía Durrieu, Alan Bush, Alicia Grande, Rikard Johansson, David Janzén, Andrea Katz, Gunnar Cedersund, Alejandro Colman-Lerner

## Abstract

Nuclear transport is an essential part of eukaryotic cell function. Several assays exist to measure the rate of this process, but not at the single-cell level. Here, we developed a fluorescent recovery after photobleaching (FRAP)- based method to determine nuclear import and export rates independently in individual live cells. To overcome the inherent noise of single-cell measurements, we performed sequential FRAPs on the same cell. We found large cell-to-cell variation in transport rates within isogenic yeast populations. Our data suggest that a main determinant of this heterogeneity may be variability in the number of nuclear pore complexes (NPCs). For passive transport, this component explained most of the variability. Actively transported proteins were influenced by variability in additional components, including general factors such as the Ran-GTP gradient as well as specific regulators of the export rate. By considering mother-daughter pairs, we showed that mitotic segregation of the transport machinery is too noisy to control cellular inheritance. Finally, we studied mother-daughter cell asymmetry in the localization of the transcription factor Ace2, which is specifically concentrated in daughter cell nuclei. We found that this asymmetry is the outcome of a higher ratio of import rate to export rate in daughters. Interestingly, rather than reduced export in the daughter cell, as previously hypothesized, rates of both import and export are faster in daughter cells than in mother cells, but the magnitude of increase is greater for import. These results shed light into cell-to-cell variation in cellular dynamics and its sources.

## Introduction

Isogenic cells, grown under identical conditions, display heterogeneity in their molecular contents. This leads to variation in their physiological states, potentially causing different cellular responses and cell fate choices. Variability emerges partly as a result of the stochastic nature of chemical reactions -specially transcription- (Elowitz et al., 2002; Raser and O’Shea, 2004; Rosenfeld et al., 2002), but also from fluctuations in other processes, such as the mitotic segregation of materials (Huh and Paulsson, 2011). In eukaryotes, preexisting differences among cells, such as in protein expression capacity, are the main sources of variation (Blake et al., 2003; Colman-Lerner et al., 2005; Gándara et al., 2019; Mitchell et al., 2016; Raser and O’Shea, 2004; Roux et al., 2015; Spencer et al., 2009; St-Pierre and Endy, 2008). These variable non-genetic traits may be passed to their descendants as cells divide, sometimes for several generations, impacting cell function (Kaufmann et al., 2007; Li et al., 2020; Mitchell, 2021; Mura et al., 2019; Spencer et al., 2009; Vashistha et al., 2021).

Regulation of nuclear transport influences important cellular processes, such as gene expression regulation and signal transduction. Despite its centrality, intrapopulation variability in transport rates has not been studied so far. Molecules move between the nucleus and the cytosol through 66 MDa nuclear pore complexes (NPCs) embedded in the nuclear envelope, which in *Saccharomyces cerevisiae* consist of 30 different nucleoporins (Rout et al., 2000) (human NPC are 110 MDa, containing 34 different nucleoporins). Passage may occur in two modes: passive diffusion for small molecules (up to 40 kDa (Paine and Scherr, 1975) or ∼ 5 nm diameter (Mohr et al., 2009; Yang and Musser, 2006)), or facilitated translocation for small and big species (Ribbeck and Görlich, 2002). In the latter, translocation is coupled to the transport of the small GTPase Ran, enabling net movement of cargoes against their chemical gradient through dissipation of the primary gradient of Ran-GTP. Facilitated translocation requires specific interactions between the translocating species and constituents of the NPC and is therefore a highly selective and regulated process. Cell-to-cell variation in nuclear transport rates could thus arise from several sources: passive transport variation should depend only on the number and quality of nuclear pores, while active transport could be affected additionally by the number of specific transporters, their activation state, the magnitude of the Ran-GTP gradient, and variability in the regulation of the affinity of the cargo for the transporters.

*S. cerevisiae* cells divide by budding, a polarized process that produces a new cell, the daughter. Transiently, daughters constitute a different cell type, in part as a result of two asymmetric (mother- or daughter-specific) genetic programs. One program drives HO expression in mothers (Bobola et al., 1996; Nasmyth, 1983; Sil and Herskowitz, 1996) while in the other program the transcription factor Ace2 induces in the early G1 of daughter cells the expression of a group of genes, including chitinase, involved in cell separation (Colman-Lerner et al., 2001). Although the ultimate mechanism that causes Ace2 asymmetric activity is not completely defined, the control of Ace2 localization has been extensively studied. Ace2 subcellular localization is cell-cycle controlled, depending on a layered regulation of its nuclear import and export. Although constantly shuttling in and out of the nucleus, from late G1 until the end of metaphase Ace2 is mostly cytoplasmic (Mazanka et al., 2008; Weiss et al., 2002). Net nuclear entry at the beginning of anaphase (Boettcher et al., 2012; Colman-Lerner et al., 2001; Mazanka et al., 2008) (note that, as in all fungi, mitosis in yeast does not involve nuclear envelope breakdown (Cavalier-Smith, 2010)), is triggered when the phosphatase Cdc14, released from the nucleolus by the mitotic exit network (O’Conallain et al., 1999), removes the Cdc28-dependent phosphorylations on Ace2. Cdc14 additionally activates the kinase Cbk1 (Brace et al., 2011). Active Cbk1 enhances Ace2 nuclear enrichment by phosphorylating it near its nuclear export signal (NES), impairing its interaction with the exportin Crm1 (Bourens et al., 2008; Brace et al., 2011; Mazanka et al., 2008; Mazanka and Weiss, 2010; Montpellier and Bourens M Bécam Am, 2008; Sbia et al., 2008). Even at this pre-cytokinesis stage when the nuclei are still connected, Ace2 appears preferentially enriched in the nascent daughter’s nucleus (Colman-Lerner et al., 2001). This asymmetry is perhaps helped by the separation of the mother-daughter nucleoplasms due to the geometric constrains of dividing nuclei (Boettcher et al., 2012). After cytokinesis, Ace2 leaves the mother nucleus but stays in that of the daughter until mid G1 (Boettcher et al., 2012; Colman-Lerner et al., 2001; Dohrmann et al., 1992; Herrero et al., 2020; Mazanka et al., 2008; O’Conallain et al., 1999; Sbia et al., 2008; Weiss et al., 2002). It is in this period in which Ace2 induces its target genes. Thus, the current view is that the asymmetric localization of Ace2 derives from inhibited nuclear export in daughter cells (Boettcher and Barral, 2014).

Herein, we developed a method to monitor nuclear transport rates in single cells using the optical technique fluorescence recovery after photobleaching (FRAP) (Axelrod et al., 1976; Cole et al., 1996b). We used this method to obtain both nuclear import and export rates in single live yeast cells. We applied it to the study of three proteins: as a passively diffusing molecule, we used yellow fluorescent protein (YFP); as actively-transported molecules, we used YFP fusions to Ace2 as well as Fus3, a mitogen activated protein (MAP) kinase in the yeast pheromone response pathway. The obtained measurements were used to determine the degree, sources, and heritability of cell-to-cell variation both in the structural components of the nuclear pore as in their function, reflected in their nuclear transport rates. Additionally, these results provide valuable insights into the mechanism behind Ace2 asymmetric localization.

## Results

### A train of FRAPs enables robust estimation of nuclear transport rates in single cells

Transport between cellular compartments *in vivo* is frequently studied using FRAP (Cole et al., 1996a; Goodwin and Kenworthy, 2005; Wachsmuth et al., 2003). In it, a perturbation is introduced by photobleaching a small area of a cell, usually with a laser. Then, this region is monitored as unbleached, bright molecules move into it, resulting in an increase (*recovery*) of the fluorescence signal. While FRAP experiments are performed in single cells, the subsequent analysis is usually done over averages of multiple cells. This is because it is difficult to obtain data in single cells with high signal-to-noise ratio (SNR) without using long exposures and/ or more intense illumination, which leads to subsequent photodamage. We reasoned that performing sequential “trains” of FRAPs on the same cell (Figure 1 A) should help improve the single-cell information (without excessive exposure or intensities), potentially avoiding the need of averaging over many cells. We assessed this possibility using a simple mathematical model (Figure 1 B) of two compartments and one shuttling species, in a fluorescent or photobleached state. We considered the photobleaching reaction to be irreversible. Fluorescent and photobleached proteins used the same import and export rates, as they are indistinguishable to the transport machinery. We simulated data consisting of trains of 1 to 5 FRAPs, using a range of realistic kinetic parameters. We introduced different levels of additive white noise to the simulated data, as it is the sort of noise expected from microscopy experiments (Gordon et al., 2007). Then, to test whether the transport rates could be estimated reliably, we analyzed the simulated data in the same way as the experimental data (described below). For very high-quality data, such as with a SNR of 50 or more, we recovered the exact rates from single FRAP experiments (mean fitted export rate= 0.92 sec^-1^ SD=0.05, true export rate=0.90 sec^-1^) (Figure 1 C). However, more likely scenarios with lower SNR showed problems. For a SNR of 5, when a single FRAP was performed, the estimated parameters had large dispersion and deviated from the simulated value significantly (mean fitted export rate= 1.57 sec^-1^ SD=2.37, true export rate=0.90 sec^-1^) (Figure 1 C). In this case, increasing the number of FRAPs per train improved the accuracy of the recovered parameters. Notably, trains of only 4 FRAPs resulted in parameter fits consistently close to the true value, even with SNR as low as 5 (mean fitted export rate= 1.05 sec^-1^ SD=0.30, true export rate=0.90 sec^-1^) (Fig 1C). Interestingly, the biggest improvement was obtained with the addition of the second FRAP on the train (Fig. 1 C). Altogether, our theoretical analysis showed that performing trains of 4 FRAPs would suffice to determine nuclear import and export rates in single cells, even for noisy single-cell data.

**Figure 1.**
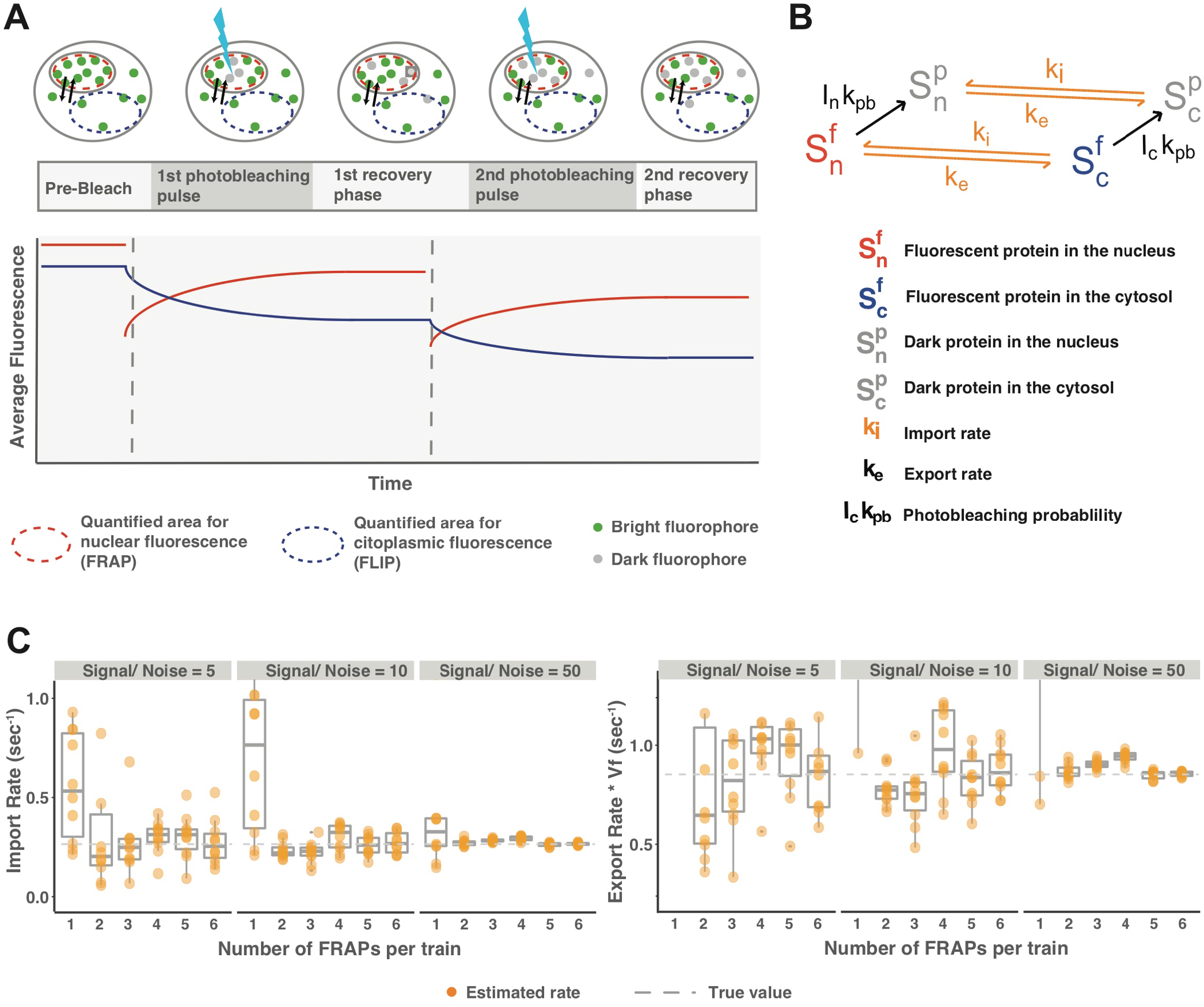
Trains of FRAPs allows the estimation of nuclear import and export rates in single cells even under low signal-to-noise conditions. *(A)* Schema of an essay with a train of 2 FRAPs. *(B)* Reactions schematic in the model used for data simulation and fitting. *(C)* Box plots show import (left) and export (right) obtained by fitting simulated data (K_I_= 0.4 sec^-1^, K_EV_= 0.9 sec^-1^) of trains of different number of FRAPs to which noise was added to achieve the signal to noise indicated. Data was simulated with K_I_= 0.4 sec^-1^, K_EV_= 0.9 sec^-1^, gray dotted line (same qualitative results were observed with other parameters). Notice that 4 FRAPs (but not 1), allow consistent recovery simulated parameters, even at realistic noise levels.

### Determination of nuclear transport rates in single cells

Based on the above analysis, we developed an experimental protocol and analysis pipeline, which we termed scFRAP (single cell FRAP), to estimate nuclear export and import rates independently for single cells (see the Appendix) (Figure 2A). Briefly, we performed trains of 4 FRAPs in yeast cells expressing either free YFP or a YFP fusion protein. We fitted the obtained data to an ordinary differential equations model that is an extension of the one described in Figure 1B (Figure 2B), in which we included the possibility that a fraction of the species be “fixed” (not shuttling) in the timescale of the experiment (e.g., molecules bound to the DNA in the nucleus).

**Figure 2.**
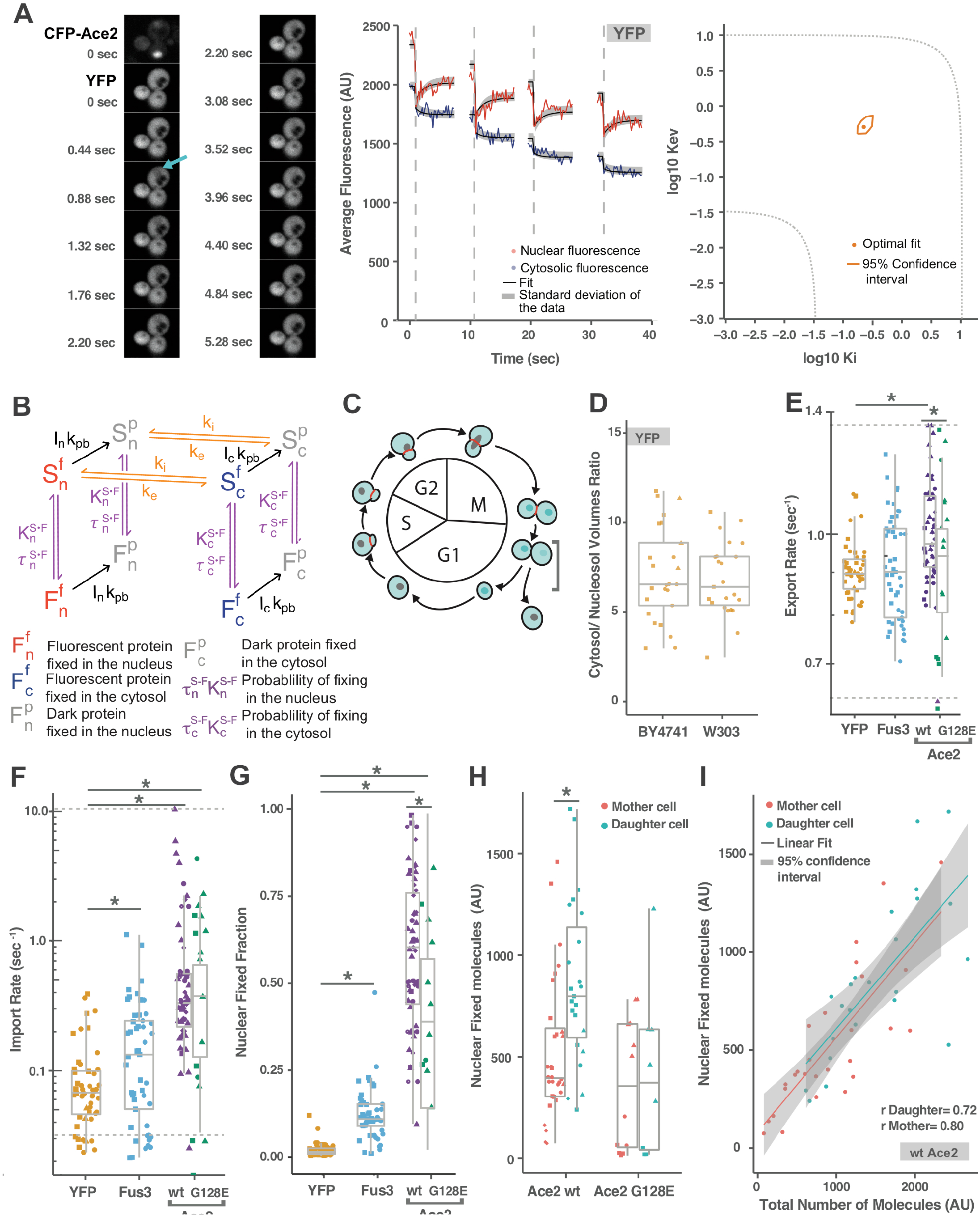
Experimental determination of nuclear transport rates in single cells. *(A)* Example of confocal-microscopy images from a representative FRAP experiment. **Left:** Time lapse images of a mother-daughter cell pair expressing Ace2-CFP and free YFP. A single image of Ace2-CFP (top left) was acquired to assess the cell-cycle stage. Then, the FRAP experiment was performed on YFP. Only the first FRAP in a train of four is displayed. After 5 initial pictures, the nucleus was photobleached (cyan arrow), then 30 images were taken to follow the recovery. Pictures were taken every 0.22 sec, but only one of two are shown. **Middle:** Fluorescence intensity curves (red: nucleus, blue: cytoplasm) and fittings (black). The dotted gray lines signal the times of the photobleaching events. **Right:** Likelihood profile analysis (see Methods). Plot shows log 10 K_EV_ vs log10 K_I_. The orange dot corresponds to the best fit and the line enclosing it delimits the 95% confidence interval (i.e. all acceptable values). Note that a closed area indicates that both rates can be identified independently. The dotted lines show the boundaries of parameter values that can be estimated using our experimental protocol (limited by the time resolution and duration of the experiment). *(B)* Diagram of the reactions included in the full model used for fitting the experimental data. In comparison to the model in 1B, we added fixed molecules (F) in the cytosol (F_c_) or nucleus (F_c_)was added. These fixed fractions can be photobleached with the same probability than the shuttling fractions. *(C)* Diagram of the yeast cell cycle. The localization of the Ace2 protein is depicted in cyan, and the actomyosin ring separating the cells, visualized trough Myo1-mCherry, is depicted in red. Yeast in the desired cell cycle-state for the experiments were chosen based on the morphological, Ace2-CFP and Myo1 information, and are marked with a square bracket. *(D)* Ratio of cytosol/nucleosol diffusion-available volumes, estimated assuming K_EV_ = K_I_ for freely diffusible YFP. The measurements were performed in two yeast strains with different background (S288C, n=25, and W303, n=23) and showed no statically significant differences (t test, p-value=0.66). *(E-F)* Estimated nuclear transport rates for the proteins YFP (n=58), Fus3-YFP (n=48) and Ace2-YFP (n=49) and Ace2 G128E-YFP (n=20). Export rates between WT and G128E-mutant Ace2 are significantly different (t test, p-value=0.04), but import rates are not (t test, p-value=0.15). *(G)* Fraction of the total amount of proteins that is fixed in the nucleus (not shuttling between nucleus and cytosol). T-test for differences between WT and G128E Ace2 p-value=0.002. *(H)* Number of nuclear fixed Ace2 molecules in mother and daughter cells, for WT and G128E mutant strains. For Ace2 WT, daughter cells have more fixed molecules than mothers (t-test, p-value=0.003). In the G128E mutant no differences were observed (t-test, p-value=0.748). *(I)* Plot shows fixed vs total Ace2. r is the correlation coefficient. Slope and intercept are not statistically different between daughters and mothers. The points represent individual cells. Distinct symbol shapes indicate experiment replicates.

In order to minimize potential variation in the transport rates due to the cell cycle, we restricted our measurements to cells at the beginning of G1 by assessing the localization of two proteins: Myo1-mCherry, a component of the actomyosin ring in the bud neck that disappears upon completion of cytokinesis (Lippincott and Li, 1998), and CFP-Ace2, which translocates to the nucleus at the end of mitosis and remains nuclear during early G1. As explained above, Ace2 remains nuclear for a longer period in daughter cells than in mothers (Colman-Lerner et al., 2001) (Figure 2C). Thus, we used cells that did not have Myo1-mCherry at the bud-neck while at the same time had nuclear CFP-Ace2 in both mothers and daughters (Figure 2C, grey bracket). Choosing this moment in the cell cycle enabled us to measure nuclear transport of Ace2 in mothers and daughters, in order to shed light on the nature of its asymmetric behavior.

Fitting of the FRAP data yielded parameters K_I_ and K_EV_. The first represents the import rate, while the latter corresponds to the export rate multiplied by **V**, the ratio of the diffusion-accessible volumes of the cytoplasm and the nucleus. Thus, to determine transport rates, we needed to determine **V**. The value of **V** is of interest by itself, and it should not be confused with the ratio of the cytoplasm to the nucleus geometric volume, since not all of the cytoplasmic and nuclear volumes are available for diffusion, due to the presence of organelles, macromolecules, chromatin and other cellular materials precluding diffusion of the protein of interest. We reasoned that **V** for each cell can be obtained for a molecule with identical import and export rates, such as YFP, which is small enough to diffuse passively through the nuclear pores, and it does not interact with any importin or exportin. Therefore, we first measured transport of YFP. To do that, we used a haploid BY4741 yeast strain (S288C genetic background) expressing YFP controlled by the *ACT1* promoter (P_ACT1_). After estimation of the nuclear transport parameters K_EV_ and K_I_ we calculated the K_EV_ to K_I_ ratio and obtained a **V** of 7.14 ± 0.53 (mean ± SEM) with an SD of 2.64 (Figure 2D). We performed the same measurement in another commonly used genetic background, W303, and obtained similar values **V** of 6.84 ± 0.43 with an SD of 2.08 (Figure 2D). This yielded import and export rates for YFP with a median value of 0.06 ± 0.02 sec^-1^ with and SD of 0.073 sec^-1^ (Figure 2 E-F, Table 1). To estimate rates for other proteins, we used the average value of **V** calculated from the YFP measurements. Doing so introduced a small uncertainty in the determination of the export rate.

**Table 1.** Estimated parameters from the scFRAP experiments. To assess the statistical significance of the estimated parameters Export rate (K_EV_), Import Rate (K_I_), tau and Nuclear fixed fraction, a Welch two sample t-test was performed on the log-transformed values. All proteins were compared to YFP (BY4741), but only significant scores are indicated (latin letters). a: P = 5.8 e-4, e: P = 1.8 e-2, f: P = 7.5 e-11, g: P = 4.2 e-06, k: P = 1.7 e-07, l: P = 4.3 e-2, p: P = 1.1 e-11, q: P < 2.2e-16, r: P = 8.6 e-06. Ace2 G128E mutant was additionally compared with Ace2 WT, and significant differences are indicated with greek letters. ρ: P = 4.4 e-2, ϕ: P = 8.0 e-3. The statistical significance of the differences in cell-to-cell variation was evaluated by an F-test to compare the variances of the log-transformed values. b: P = 4.5 e-3, c: P = 3.8 e-3, d: P = 1.6 e-3, h: P = 3.2 e-2, i: P = 1.0 e-2, j: P = 2.4 e-2, m: P = 5.1 e-3, n: P = 4.9 e-3, o: P = 8.5 e-3, s: P = 1.3 e-05, t: P =3.4e-07, u: P = < 2.2e-16, v: P =< 2.2e-16.

| Protein | Export Rate<br>(median ± se)<br>[sec<sup>-1</sup>] | CV<br>Export<br>Rate | Import Rate<br>(median ± se)<br>[sec<sup>-1</sup>] | CV<br>Import<br>Rate | Tau<br>(median ± se)<br>[sec] | CV Tau | Nuclear<br>Fixed<br>Fraction<br>(median ± se)<br>[sec<sup>-1</sup>] | CV Nuclear<br>Fixed<br>Fraction |
| --- | --- | --- | --- | --- | --- | --- | --- | --- |
| YFP<br>(BY4741) | 0.06 ± 0.01 | 0.87 | 0.07 ± 0.01 | 0.89 | 2.10 ± 0.25 | 0.54 | 0.01 ± 0.01 | 1.25 |
| YFP<br>(W303) | 0.06 ± 0.02 | 0.86 | 0.07 ± 0.02 | 0.89 | 2.02 ± 0.21 | 0.51 | 0.01 ± 0.00 | 0.59 s |
| Fus3<br>(W303) | 0.06 ± 0.02 | 1.16 b | 0.13 ± 0.03 e | 1.12 h | 1.72 ± 0.36 | 0.94 m | 0.11 ± 0.01 p | 0.64 t |
| Ace2<br>(BY4741) | 0.11 ± 0.04 a | 1.30 c | 0.33 ± 0.24 f | 1.85 i | 0.93 ± 0.35 k | 2.07 n | 0.61 ± 0.03 q | 0.37 u |
| Ace2<br>G128E<br>(BY4741) | 0.08 ± 0.04 $\rho$ | 1.26 d | 0.37 ± 0.11 g | 1.06 j | 0.88 ± 0.42 l | 0.99 o | 0.39 ± 0.06 r,<br>$\phi$ | 0.70 v |

We applied this method to study the nuclear transport of Ace2 using a strain expressing *P*_*ACE2*_*-YFP-ACE2*. For comparison, we additionally measured in a strain expressing *P*_*FUS3*_*-FUS3-YFP*, coding for an unrelated protein kinase Fus3, central in the pheromone response pathway, which also shuttles between the nucleus and the cytoplasm (Dohlman and Thorner, 2001). When fused to YFP, these proteins have a size of ∼115 kDa and ∼69 kDa, respectively. The former is too large to shuttle by passive diffusion through the NPCs, while the latter is too close to the size limit to be certain (Paine and Scherr, 1975). In fact, a previous FRAP-based study showed that a Fus3-GFP fusion shuttles both by facilitated transport and to a lesser extent by passive diffusion (van Drogen et al., 2001).

The export rate for Fus3-YFP did not differ from that of YFP, but Fus3 import was faster (Figure 2 E-F, Table 1). In contrast, import and export of YFP-Ace2 was faster than YFP, (Figure 2 E-F, Table 1). We also measured the behavior of the mutant Ace2-G128E (Racki et al., 2000). This mutation lies within Ace2 NES region (amino acids 122-150) (Jensen et al., 2000), and shows an altered cellular localization, persisting in the mother cell nucleus to mid G1 phase, similar to daughter cells (Colman-Lerner et al., 2001). Mutations in the NES region impair the interaction with the exportin Crm1 (Bourens et al., 2008; Mazanka et al., 2008). In agreement with these results, our method revealed that export of Ace2-G128E was slower, while its import rate was unchanged (Figure 2 E-F, Table 1). Taken together, these results support the notion that our method is suitable for estimating import and export rates of endogenous proteins in single yeast cells with enough precision to detect functional differences.

### Estimation of the amount of non-shutting molecules in the nucleus

The FRAP-analysis method also renders an estimation of the number of non-shuttling (“fixed”) molecules present during the experiment in either compartment (Figure 2B). Note that these molecules are only “fixed” in the time scale of the experiment (seconds), but they could be binding and unbinding from their “traps” in a slower timescale. Our analysis suggested that for the proteins studied here there were no fixed molecules in the cytoplasm (see Supplementary Material). We found no fixed molecules in the nucleus for the case of YFP (Figure 2G), as expected. In contrast, of the nuclear pool, on average, about 10% of Fus3-YFP and 60% of YFP-Ace2 molecules were fixed (Figure 2G). The MAPK Fus3 targets several DNA-associated substrates in the nucleus, such as the transcription factor Ste12 and its regulators Dig1 and Dig2 (Dohlman and Thorner, 2001). Thus, this 10% of fixed molecules could represent Fus3 bound to these species, which themselves do not shuttle (van Drogen et al., 2001). In the case of Ace2, since it is a transcription factor, the high amount of fixed Ace2 molecules likely corresponded to DNA-bound Ace2. Notably, daughter cells had more fixed Ace2 than mothers (Figure 2H), consistent with Ace2 preferential induction of genes in daughter cells (Colman-Lerner et al., 2001). Single cell analysis revealed that the number of fixed Ace2 molecules in nuclei correlated well with the total number of Ace2 molecules in a cell, independently of cell-type (Figure 2I, SM Figure 1). This suggested that there was not a particular enhancement of Ace2 DNA binding in daughters, there was just more Ace2 in them. In the Ace2-G128E mutant, which exhibits symmetric localization and expression of its target genes (Colman-Lerner et al., 2001), mother cells had a similar number of nuclear fixed YFP-Ace2 molecules than daughters (Figure 2H, SM Figure 1). Finally, the number of nuclear fixed molecules did not correlate with the import/export rates ratio (SM Figure 1).

### The high variability in nuclear transport rates is mainly due to variation in the amount of NPCs

Interestingly, we found large cell-to-cell variation in the import and export rates of all three proteins studied (Figure 2 E-F, Table 1). This variability was significantly higher for YFP-Ace2 and Fus3-YFP than for YFP. The high cell-to-cell variation was evident in the raw traces, and thus we do not believe it might be the result of misfitting (Figure 3A).

**Figure 3.**
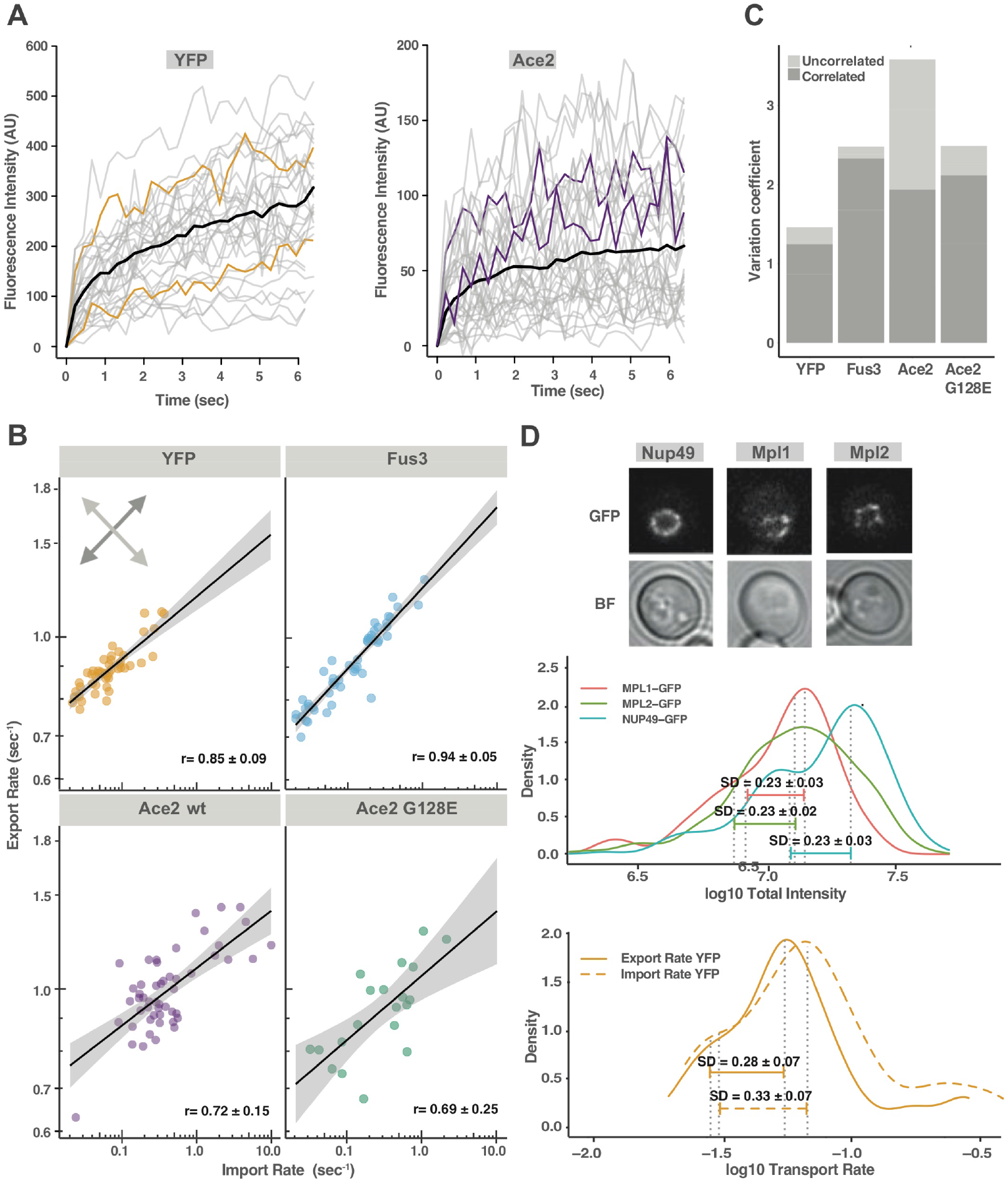
The high cell-to-cell variation in nuclear transport rates is mostly due to variation of general transport machinery. (*A*) Individual traces of FRAP experiments in single cells (gray lines). Each line corresponds to the average of the four FRAPs in the train where the first value after photobleaching has been subtracted. The average of all traces is shown in black. Representative fast and slow traces are highlighted in color for both proteins. (*B*) Plots show export vs import for the proteins YFP (n=58), Fus3-YFP (n=48) and Ace2-YFP (n=49) and Ace2 G128E-YFP (n=20). The black line corresponds to linear model, and the gray shadowed area represents de 95% confidence interval. r values correspond to the correlation coefficient. Errors were calculated by bootstrapping and correspond to 95% confidence interval. (*C*) Plot shows a quantification of the correlated and un-correlated variation between import and export rates. The correlated components represent the spread of the normalized data on the direction parallel to the linear fit (dark gray arrow on B). The un-correlated component is a measure of the normalized rates perpendicular to the linear fit (light gray arrow in B). The details of the estimators can be found in the STAR methods section. Correlated variation could be attributed to heterogeneities in general transport machinery, such as NPCs. (D) Histograms (in the form of density plots) show the frequency of the log transformed of values. Color segments represent the SD of the log-transformed values and can be used to compare the variability. The errors in the SD were calculated by bootstrapping and correspond to 95% confidence interval. **Top:** Representative confocal images of the analyzed nuclear pore proteins. **Middle:** Total fluorescence of GFP fusions to the indicated nuclear envelope proteins per cell, as quantified from fluorescence intensity (AU). MPL1 (NPC associated, n=2254), MPL2 (NPC associated, n=1967) and NUP49 (NPC integral, n=1442). **Bottom:** Density plots as above for Import Rate (sec^-1^) and Export rate (sec^-1^) of free YFP. Note that the variability in the NPC components MPL1, MPL2 and NUP49 is similar than for the nuclear transport rates of all proteins analyzed.

We then wondered what the sources of the measured variability in rates were. We reasoned that some of it might be due to cell-to-cell differences in the abundance or the activity of components of the core nuclear transport machinery (i.e., those that affect equally import and export, such as NPCs). Variation in the rates of actively transported proteins, however, could additionally be affected by cell-to-cell variability in the abundance of nuclear transporters, regulators, and the Ran-GTP gradient. Importantly, variability in core nuclear transport machinery and the Ran-GTP gradient would impact import and export rates in the same manner, increasing their correlation, while variability in the regulators could affect these rates to different extents, decreasing their correlation.

Thus, to dissect these two sources of variation, we quantified the correlated and uncorrelated components of the variability in the rates of each protein (Qiuyan Fua and Pachter, 2016) (Figure 3 C). For the three proteins studied, we found that import and export rates were highly correlated (Figure 3B, C). In the case of YFP, this correlated component represented almost all the variation. This result was expected, since YFP passage through the NPC does not require binding to any factors, and thus cell-to-cell variation is most likely dominated by differences in the number or quality of NPCs (see below). Although more variable than YFP, Fus3 maintained the high import to export rates correlation, indicating that whatever additional factors contribute to the variation, they affect nuclear passage in both directions. Considering that Fus3 transport is at least partially active, we speculated that a relevant factor is variability in the strength of the Ran-GTP gradient. In contrast, for Ace2, we found a greater amount of uncorrelated variability (Figure 3 B and C), probably because its import and export are individually regulated. Remarkably, Ace2 uncorrelated variability was greatly reduced in the mutant Ace2-G128E (Figure 3C)). Since the G128E mutation reduces Ace2 interaction with exportin Crm1 and reduces the effective export rate (Figure 2E), the high Ace2 uncorrelated variation could be attributed to cell-to-cell variability in the export rate. Heterogeneity in Ace2 export could originate in part in mother-daughter differences, but also in the strong cell cycle regulation of its nuclear localization. Despite the fact that we only imaged cells that were traversing a specific, and short, part of the cell cycle (just post cytokinesis) small differences in their actual cell cycle position were unavoidable, which might contribute to variability in the measured transport rates.

To assess whether the co-variation of import and export rates is due to differences among cells in the number of NPCs, we analyzed, as a proxy, the cell-to-cell variability in the amounts of nuclear pore proteins (Khmelinskii et al., 2010). We quantified nuclear envelope fluorescence in confocal images of yeast cells expressing three different GFP-tagged proteins: NPC-associated proteins Mlp1 and Mlp2, and the central core component of the NPC Nup49 (Huh et al., 2003) (Figure 3 D, top). Interestingly, the magnitude of the cell-to-cell variation in these nuclear pore proteins is similar to that of nuclear transport rates (Figure 3D). This suggested that the variability in YFP nuclear transport rates may be directly attributable to variability in NPC number. If so, our results suggest that other possible sources of variation, such as variable amounts of regulators or the quality of the nuclear pores being heterogenous, have a relatively small contribution to overall variation in transport rates.

### Segregation of NPCs, and nuclear transport rates during cell division are highly noisy

A possible mechanism creating population variability in NPC abundance, which in turn might translate into variation in transport rates, is their random segregation during cell division. To explore this effect, we asked if the amount of NPC proteins was correlated between mother cells and their daughters. (Note that since our measurements were performed just after cell division, it was warranted to assume that there was no substantial contribution of newly synthesized NPC components.) If so, that would indicate that daughters receive a consistent fraction of the NPCs with low segregation noise. Comparison of mother vs daughter NPC numbers revealed no significant correlation (Figure 4 A), suggesting that segregation for NPCs is highly noisy. Similarly, nuclear transport rates of YFP did not correlate between mother and daughter cells, nor did those for the actively transported proteins Fus3-YFP and YFP-Ace2 (Figure 4 B-C). We confirmed all the above results using a similarity measure (SM Figure 2 A). These findings show that, even in the first generation, there is no epigenetic inheritance of NPC abundance or nuclear transport rates.

**Figure 4.**
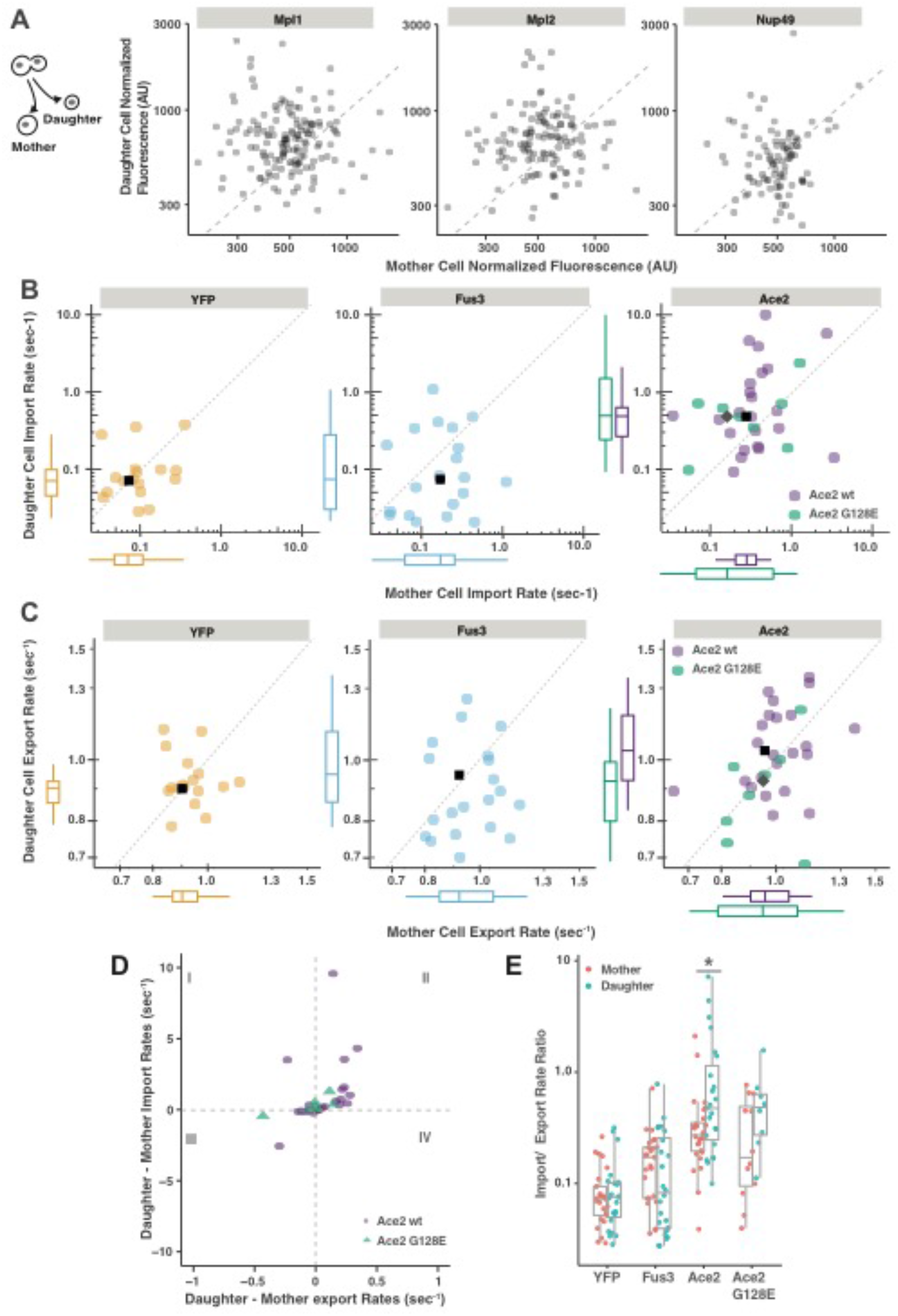
Mother- Daughter cell segregation of NPCs and transport rates. (*A*) Plots show daughter vs mother normalized nuclear fluorescence (total GFP fluorescence in the nucleus divided by the cell volume) of three NPC components, Mlp1 (left), Mlp2 (middle) and Nup49 (right). (*B-C*) Plots show daughter vs mother import rate (*B*), or export rate (*C*), obtained for YPF (left), Fus3-YFP (middle) and YFP-Ace2 (right, WT: green or the G128E mutant: purple). Transport rates of WT YFP-Ace2 were significantly different between mother and daughter cells (t-test p-value for export rate= 0.06, import rate= 0.01). (*D*) Plot of the difference in rates (daughter minus its mother), import vs export for YFP-Ace2 (WT, blue circles) and the G128E mutant (green triangles). Most WT YFP-Ace2 WT cell pairs fall in quadrant II, indicating that daughter cell has both higher import and export rates, while for the YFP-Ace2-G128E mutant most pairs are close to the origin, indicating small differences. (*E*) Ratio of import to export rate for the indicated strains in mothers (red) and daughters (green). A-C, black squares (and grey diamond in C) correspond to the location of the mean of the Y and X axes.

### Mother-Daughter asymmetry in nuclear transport rates

Another mechanism that could cause population variability would be a systematic asymmetry in the segregation of components during division. As mentioned above, yeast division is polarized, with the mother cell forming the bud that will become a daughter cell. Thus, for example, it could be that daughters receive, on average, more (or less) NPCs, or that they are of a different quality. We found no significant differences between the mother and daughter cell population in nuclear transport rates for either YFP (t-test kE p= 0.47, K_I_ p = 0.46) or Fus3-YFP (t-test kE p = 0.48, K_I_ p = 0.21) (Figure 4 B-C, left), indicating that whatever differences might exist between mothers and daughters, these are not sufficiently strong to overcome the noisy segregation of NPCs. Thus, we concluded that NPC function is, in this regard, symmetric (similar in mothers and daughters).

In stark contrast with YFP and Fus3-YFP, for YFP-Ace2 the rates of both import and export were significantly faster in daughter cells than in mothers (t-test kE p = 0.06, K_I_ p = 0.01) (Figure 4 B-C, right). In consonance with the above population-level results, we found that, when compared pair-wise, most daughter cells have both faster export and import of Ace2 than their mothers (Figure 4 D), an intriguing result that would require future studies. It is noteworthy that also the ratio of import to export rate was larger in daughters (Figure 4 E). Taken together, these results indicated that the mobile (i.e., not fixed) molecules of YFP-Ace2 shuttle faster and are more nuclear in daughter cells. Finally, this kinetic difference between mothers and their daughters was not observed in the symmetrically localized YFP-Ace2 G128E mutant (Figure 4 D-E), free YFP, or Fus3-YFP (Supplementary Figure 2 B), so it appears to be related to the specific mechanisms that cause Ace2 asymmetric localization to the daughter cell.

## Discussion

Quantitative measurements of dynamic processes in single cells are essential for a system-level understanding of cellular function. Here, we introduced a method combining a FRAP protocol with an analysis pipeline for the determination of nuclear import and export kinetic rates in intact single cells, suitable for the usually noisy experimental conditions. This method also extracts the fraction of non-shuttling molecules, a parameter of significant biological value. This method, developed for nuclear transport in yeast, may be adapted for other cellular compartments, including membrane-less organelles, as well as in other cell types, such as mammalian cells. We applied this method with two central goals: to obtain a better understanding of cell-to-cell variability in kinetic rates and to study the mechanisms behind a cell-fate decision (the asymmetry in the transcription factor Ace2 that drives a daughter-cell specific genetic program). Our estimations matched well with previous population measurements and were affected as expected by genetic manipulations.

We estimated nuclear transport rates in the order of 0.1 sec^−1^. This number means that, if there are 1000 molecules of a protein X in a yeast cytosol (approximately 1 µM), and 86 nuclear pore complexes in the nuclear envelope (Winey et al., 1997), 1.2 molecules of X are imported to the nucleus per second per pore. This agrees well with the estimations of 0.18-3.3 molecules transported per pore per second under a 1 µM gradient previously (Nemergut and Macara, 2000; Ribbeck et al., 2001; Siebrasse and Peters, 2002; Timney et al., 2006). This is remarkable, as the cited studies were performed in a different cell type, with different cargos, and under different conditions. If true, this would imply that NPC permeability has been strongly conserved, at least between fungi and mammals.

Translocation of the actively transported proteins we tested was faster than the passively diffusing YFP. Active transport rates can exceed passive diffusion, sometimes by a large factor, as established by single transporter recordings (Kiskin et al., 2003; Siebrasse and Peters, 2002) as well as by bulk import measurements in permeabilized cells (Frey et al., 2018; Ribbeck et al., 2001). The pheromone response pathway MAPK Fus3 was exported at rates similar than YFP, but it was imported twice as fast. From this we concluded that at least a sizable fraction of Fus3-YFP is actively transported. This is in good agreement with earlier findings (van Drogen et al., 2001) that blocking active transport reduced, but did not eliminate, Fus3 nuclear transport.

We found large cell-to-cell variation in transport rates for the three proteins studied. When we compared import and export rates in single cells, we found that most of this variation was correlated, with some cells having fast and some cells slow transport dynamics. Considering that YFP only crosses the NPC by passive diffusion, this finding suggested that the heterogeneity in the transport rates could be due to variability in NPC abundance, the extent of which we measured by direct labeling of NPC components. Thus, heterogeneity in the number of NPCs is likely to be the main determinant of variability in the kinetic rates.

The actively transported proteins Fus3 and Ace2 had higher levels of correlated variation than YFP. The molecular mechanism responsible could be intrinsic to Fus3 or Ace2 transport, or it might be extrinsic and as such be a more general phenomenon, impacting all actively transported proteins. A general effect could result, for example, from variability in the strength of the Ran-GTP gradient. We found no previous data investigating this possibility.

Variability in the core transport machinery (leading to the correlated variability import and export rates we showed here) would only affect the shuttling speed of mobile proteins, without affecting their net subcellular localization (which is dictated by the ratio of import to export). Whether and how shuttling speed could control protein activity has not been established. One possible scenario could arise for a protein whose nuclear transport is coupled to its post-translational modification in only one compartment and the removal of that modification in the other. If this modification proceeds at a rate slower than transport, then proteins that shuttle faster would be less available for the modifying enzyme. Naturally, variability in net import or export would be important for molecules that are transported in only one direction, such as newly synthesized RNA molecules. Cells with faster nuclear export should have mRNAs more readily available for translation, leading to a faster expression response. In the case of Fus3, one could imagine that cells with faster Fus3 shuttling might be able to mount a transcriptional response to mating pheromone faster, which might lead to faster mating. Thus, it would be interesting to further study the basis of this regulation, and to determine if MAPKs in other systems, such as mammalian cells, also have variable shuttling speed.

According to our estimations, the effective volume in the yeast cytosol available for diffusion is 7 times larger than that volume in the nucleosol. Since the yeast nucleus is approximately 14 times smaller than the cell, as determined using geometrical considerations only (Jorgensen et al., 2007), it follows that the cytoplasm is twice as occupied as the nucleus, perhaps due to the presence of organelles. Indeed, using data obtained by soft X-ray microscopy (Uchida et al., 2011), we estimated a ratio of 8 subtracting from the cytoplasm volume that of the vacuole, mitochondria, and lipid bodies. Also, our number agrees well with electron microscopy-based estimations of the cell / nucleus (subtracting organelles) volume of 6.25 (Webster et al., 2010). The agreement is remarkable since we obtained this ratio solely from the dynamics of YFP in the FRAP experiments.

Cellular division involves the segregation of all its components between descendant cells. While a noisy partitioning of cellular components would increase the population variability (Huh and Paulsson, 2011), its counter-side, “cellular heritability”, would limit it (Vashistha et al., 2021). Interestingly, our results show no heritability in either the number of NPCs or nuclear transport rates, suggesting that partitioning of the nuclear transport machinery during cell division is highly stochastic. Epigenetic heritability of cellular characteristics has been reported in several processes, where it was found to last between 2 to 10 generations, with first-generation correlations of 0.6-0.9 (Kaufmann et al., 2007; Mura et al., 2019; Sigal et al., 2006; Spencer et al., 2009; Vashistha et al., 2021). There are also reports of noisier processes where no heritability was found (Li et al., 2020; Suel et al., 2006).

Although the ultimate mechanism that causes Ace2 asymmetric localization to the daughter cell nucleus is still unknown, our FRAP measurements provide notable insights. First, consistent with reports of enhanced transcription in daughter cells, we found a higher amount of Ace2 fixed in the nucleus in this cell type, which seems a consequence of the higher concentration of Ace2 in the nucleus of daughters, not the result of additional regulation. Second, contrary to our expectations based on previous cumulative data, we did not find that reduced nuclear export in the daughter cell could explain Ace2 asymmetric enrichment in the daughter’s nucleus. Rather, our surprising results showed that, for most daughters, both import and export were faster than in their mothers. The higher nuclear enrichment in the former is the result of a higher import/export rates ratio. In the Ace2 G128E mutant, which displays symmetric Ace2 localization between mother and daughter cells, the import/ export rates ratio was also symmetric, further supporting that this ratio is responsible for the net localization of Ace2, rather than individual rates. We note that our results are biased to the early stages of the establishment of the asymmetric localization, a time window in which it is still possible to detect Ace2 in mother nuclei. It is conceivable that at a later stage, transport dynamics of Ace2 change so that export slows down in daughters and speed up in mothers.

Altogether, this study sheds light into cell-to-cell variation in the kinetic rates of nuclear transport and its sources. It is likely that a similar heterogeneity occurs in other cellular processes. High population variability in kinetic rates could have a big impact on cellular function, and future studies will be needed to understand how it propagates and the consequences it has.

## Acknowledgements

We are grateful to Peter Pryciak and Alejandra Ventura for feedback and advise. We also thank Rich Yu for providing plasmids. We are especially grateful to Linnea Järvstråt, Ulrike Münzner, Karin Wiberg, and Martin Gollvik who participated in different aspects of the modeling and parameter estimation work. Work at GC lab was funded by the Swedish Research Council and at ACL lab by grants PICT2005-33628, PICT2007-847 and PICT2010-2248 from the Argentine Agency of Research and Technology (ANPCyT) to ACL.

## Authors contributions

LD performed most of the scFRAPs, constructed the strains and some of the plasmids used in the study, performed modeling and data analysis. AB performed scFRAPs, data analysis, developed the model and the R package for data fitting. AG performed the measurements of nuclear pore proteins. AK wrote the R package for data fitting. LD, AB and ACL interpreted the data. RJ supervised DL and MG and performed the analysis with multiple shell models. DL performed the profile likelihood analysis, and the implementation with the local optimization algorithm; MG performed the analysis proving that a detailed mechanistic model is equivalent to a two-state model; GC directed the theoretical aspects of the project. LD and ACL conceived the project. LD, AB, and ACL wrote the paper.

## STAR Methods

### RESOURCE AVAILABILITY

#### Lead contact

Further information and requests for resources and reagents should be directed to and will be fulfilled by the lead contact, Alejandro Colman-Lerner.

#### Materials availability

Plasmids and yeast strains generated in this study are available upon request.

#### Data and code availability

- The datasets resulting from the quantification of the FRAP experiments study are publicly available, and can be found are listed in the key resources table as of the date of publication. The DOI is listed in the key resources table. Raw microscopy images reported in this paper will be shared by the lead contact upon request.
- The NuclearFRAP R package, has been deposited at Zenodo and is publicly available as of the date of publication. DOIs are listed in the key resources table.
- Any additional information required to reanalyze the data reported in this paper is available from the lead contact upon request.

### EXPERIMENTAL MODEL AND SUBJECT DETAILS

#### Saccharomyces cerevisiae strains and plasmids

We performed general molecular biology procedures, yeast strain manipulation and construction according to previously established methods (Ausubel et al.; Guthrie and Fink, 1991)

All strains generated and used in this study are listed in the key resources table.

Tagging of Myo1 with mCherry was performed by homologous recombination by transforming the appropriate strains with a PCR fragment containing mCherry followed by a hygromycin resistance cassette, flanked by 40 nucleotides of homology with the end of the MYO1 ORF (except the STOP codon) on its 5’ end and the region of the 3’ UTR from +50 to +90. We used as template for this PCR plasmid Pry2 2060.1 Hyg (Table 2). The sequence of the primers is listed in the the key resources table.

We selected transformants in plates with YPD plates with hygromycin and confirmed that integration occurred in the right location by visual determination of the presence of the red fluorescent ring at the bud-neck.

All plasmids generated and used in this study are listed in the key resources table.

#### Cells preparation for microscopy

Yeast cells were prepared for the scFRAPs experiments by growing them at least two consecutive days on selective BSM-TRP LEU URA 2% glucose plates. Then yeast from the least grown part of a streak were picked, resuspended in BSM-TRP, LEU, URA 2% glucose liquid media and added to a 384-well glass bottom plates (MGB101-1-1-LG, Matrical Biosciences). To prevent cells from moving, we pre-treated the wells with concanavalin A (type V; Sigma-Aldrich, St. Louis, MO). To do this, we added to each well 50 ul of a 100 mg/ml solution of concanavalin A in water, incubated at least for 20 min at room temperature (18–23 1C), and then washed 2 times with water. Cells were allowed to settle on the plate for 10-20 min and then we removed the supernatant and added 50 µl of fresh selective medium.

### METHOD DETAILS

#### Imaging and scFRAPs assays

The scFRAPs were performed in an Olimpus IX-81 inverted microscope with a FV1000 confocal module with an oil immersion Olympus UplanSapo 63X objective (numerical aperture, NA 1.35). For the experiments we used an automatic z-axis control, a motorized x-y stage, a 458-488-515 argon and a 543 He-Ne lasers, and Hamamatsu R6353 photomultipliers (PMTs).

Cells in the desired stage of the cell cycle were identified, then reference images of the transmission, CFP and mCherry channels (to observe Ace2-CFP and Myo1-mCherry) were acquired. Then 4 consecutive FRAPs were performed in the YFP channel. Each FRAP consisted of 5 initial images, followed by a partial photobleaching of the nucleus, and 30 images to follow the recovery. The time resolution was 0.2 sec.

#### Data Analysis and Fitting

For a detailed description of the protocols for image acquisition, data correction and fitting, see the Appendix. Briefly, the images were quantified with the ImageJ (Rasband, 1997-2005) plugin Time Series Analyzer. Then, model fitting was performed in R (Team, 2010) with our own R packages, Rcell (Bush et al., 2012) and NuclearFRAP.

### QUANTIFICATION AND STATISTICAL ANALYSIS

#### Statistical testing

The number of data points -individual cells or mother-daughter pairs of cells depending on the case-for each analysis can be found in the figure legend.

All estimated parameters are presented as median ± S.E.M. When indicated, statistical significance was calculated using Student’s two-sample t-test (unpaired two samples for means) over the logarithms of the values (with base 10). P-values are informed only for significant (p<0.05) differences.

Comparisons between variances were performed according to (Lewontin, 1966). The variance of the logarithms of the values (with base 10) was contrasted by an F-test. P-values are informed only for significant (p<0.05) differences. This analysis permits comparison of relative variances for values with different means, without requiring normality.

The error bars for the correlation and SD in Figure 3 were produced by bootstrapping 1000 times with reposition samples of 90% the size of the data for the experiment, estimating either the correlation or the SD according to the case, and then defining the interval that contained 95% of the estimated values.

#### Estimation of correlated and un-correlated variability

For the analysis in Figure 3, where the cell-to-cell variability in transport rates was decomposed into correlated and un-correlated, the following estimators were used (after (Qiuyan Fua and Pachter, 2016)).

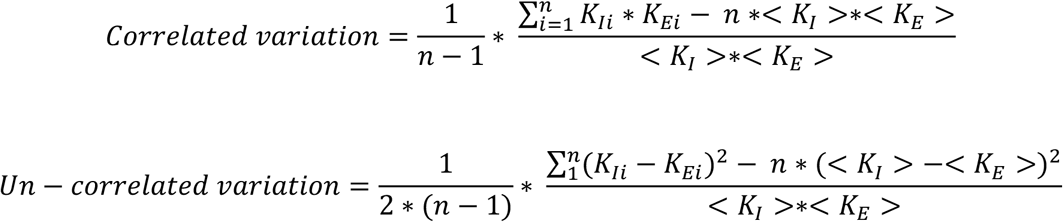

#### Data inclusion/ exclusion decision

Prior to analysis and fitting, the acquired images were manually curated in order to discard experiments that had technical issues, such as cells movement or focus drift. After fitting, data for which no statistically acceptable parameters were found was filtered out. This represented approximately 15% of the single-cells data, and was due in general to the optimal parameters laying outside of the pre-defined region of the parameter space that our experimental protocol could reliably assay. No data points were excluded during the biological analysis.

---

**Supplementary Figure 1 (related to Figure 2).**
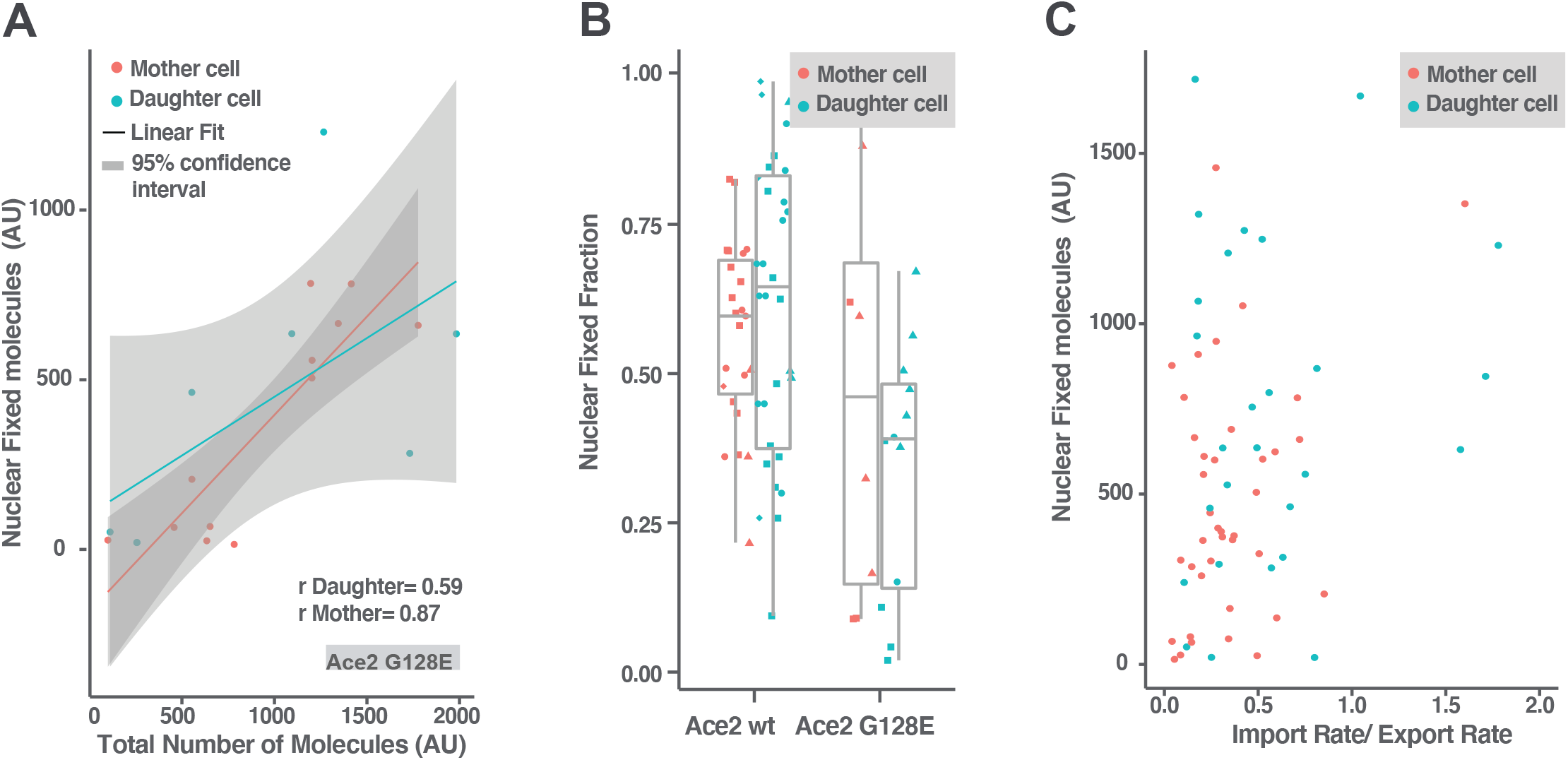
Mother- daughter cell asymmetry in the distribution of Ace2 mobile and fixed molecules. (A) Plot shows fixed vs total Ace2-G128E. r is the correlation coefficient. Slope and intercept are not statistically different between daughters and mothers. The points represent individual cells. The symbol shapes indicate experiment replica. Note that the Ace2 WT data is in Fig. 2 I. (B) Fraction of nuclear fixed Ace2 molecules in mother and daughter cells, for wt and G128E mutant strains. For both Ace2 wt and G128E mutant we found no significant differences. (C) Plot shows amount of Ace2 WT fixed molecules in the nucleus vs the import/export rates ratio for mother and daughter cells, which reflects the fraction of the mobile molecules that are nuclear. We found no significant correlation.

**Supplementary Figure 2 (related to Figure 4).**
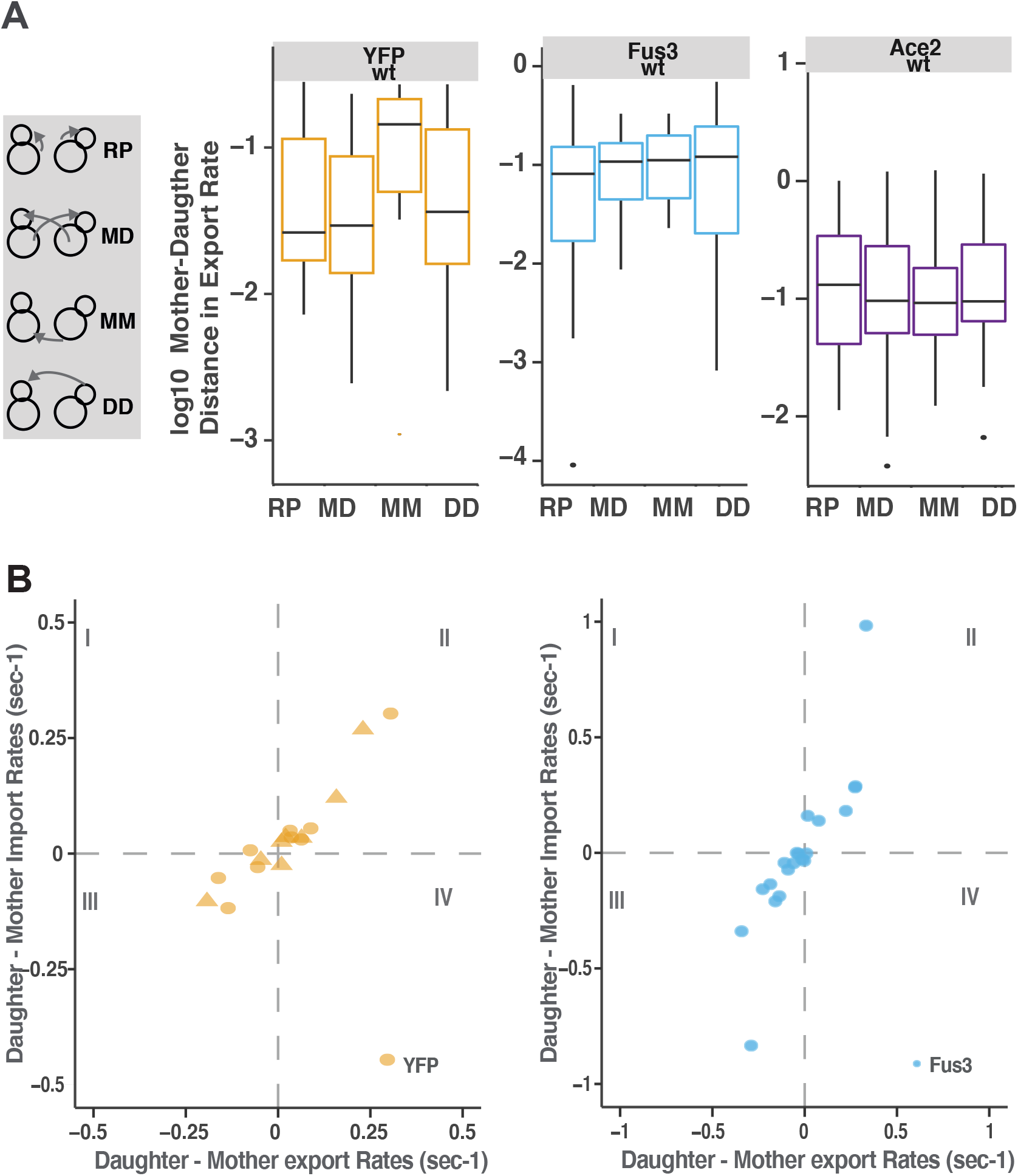
Difference in nuclear transport rates between mother and daughter cells. (A) Similarity between export rate in pairs of cells in the case of YFP (left), Fus3-YFP (middle) and YFP-Ace2 (left). Plots show distance (the square of the difference in rates) in export rates between the mother and the daughter cells. Pairs of cells were generated for comparison by bootstrapping with either 2 random cells (RP), two random mothers (MM), two random daughters (DD), or the real mother-daughter pair (MD). All the groups have the same size as the MD pairs (n YFP= 17, n Fus3=20, n Ace2 WT= 23). (B) Plots show the difference in rates (daughter minus its mother), import vs export for YFP (left) and Fus3-YFP (right).

