## Supplementary Material for "Characterization of cell-to-cell variation in nuclear transport rates and identification of its sources"

### Appendix for

### High cell-to-cell variability in nuclear import and export rates

#### 1. Modeling a fluorescence recovery after photobleaching (FRAP) experiment

The raw data obtained from any FRAP experiment only indicates whether the protein is shuttling in and out of the bleached region. To extract kinetic rates, we fitted a mathematical model to the experimental data. For this purpose, we wrote an ordinary differential equations (ODEs) model in which the inward and outward fluxes depend linearly on the nuclear and cytosolic fluorescent protein concentrations. In the model, n is the number of fluorescent molecules in the nucleus, c is the number of fluorescent molecules in the cytosol, $k_I$ is the import rate and $k_{EV}$ is the export rate multiplied by the ratio of cytoplasmic and nuclear accessible volumes (the volume through which the molecules can diffuse). With these notations, the model is given by

$$\frac{dn}{dt} = -n\, k_{EV} + c\, k_I \qquad (1a)$$

$$\frac{dc}{dt} = n\, k_{EV} - c\, k_I \qquad (1b)$$

The model (eq 1a and 1b) depends on a number of assumptions. Firstly, it is assumed that the fluxes before the photobleaching event are at steady-state. Interestingly, this steady-state assumption means that linear kinetics follows by necessity (see section 1.1). Another assumption behind the model is that the cytosol and the nucleus may be considered as well-stirred compartments. This is supported by three arguments. The first argument is based on a test done by a dedicated FRAP in which we photobleached completely a small area (~0.5-micron diameter) within a given compartment of a cell expressing YFP and then imaging the cell to determine how fast the bleached area recovered its original fluorescence level. Both in the nucleus and in the cytosol this time was less than 1 second. Since it takes at least 5 seconds for the YFP signal to reach steady state when the whole nucleus is photobleached (Appendix Figure 2A), we conclude that diffusion is much faster within than between compartments. The second argument is based on a theoretical approach, where we estimated the time scale for diffusion by dimensional analysis. We calculated the time that it would take a protein to diffuse a given distance “r” as the ratio between $r^2$ and the diffusion coefficient for that protein. For a small protein like YFP (27 kDa), this calculation gives 5 ms to diffuse 2.5 µm (half the radius of an average yeast cell); for a bigger protein of ~86.5 kDa) it would take 50ms, still significantly faster than the estimated transport between compartments. The third argument is based on a comparison of other model-structures, where the cytosol was divided in 2 to 5 concentric shell-like compartments (like layers of an onion) (see section 1.2). For none of these models we found a significant improvement on the fits compared to the simpler model (Appendix Figure 1). Thus, taken together, the above arguments suggest that the main assumptions behind the model are reasonable.

##### 1.1. A linear model follows from the assumption of initial steady-state.

Let N be the total number of molecules in the cell, both fluorescent and bleached, of our protein of interest. Let $N_c$ denote the cytosolic fraction, and $N_n$ the nuclear fraction. At steady state these fractions remain constant so that the flux of molecules leaving the cytosol and entering the nucleus, $R_I$, is constant in time and exactly as big as the flux of molecules leaving the nucleus, $R_E$. Assume that the system is in such a steady-state.

Let the superscripts **b** and **f** denote the fractions of N that are bleached and fluorescent, respectively. In other words, $R^b_I$ denotes the flux of bleached molecules that are entering the nucleus. Since a molecule is either bleached or fluorescent, the flux in the direction x, $R_X$, is the sum of the flux for the bleached and fluorescent fractions respectively, $R_x = R^b_x + R^f_x$.

Assume that the bleached and fluorescent molecules are equal in all matters except detection, e.g. that they are homogenously mixed in the compartment they occupy, that they have equal affinity for the transporter, whether passive or active, etc. With this assumption it follows that a specific fluorescent molecule in the cytosol is, on average, equally likely to enter the nucleus as another specific non-fluorescent molecule in the cytosol. In other words, the fraction of the fluorescent and non-fluorescent fluxes will be exactly given by the fraction of fluorescent and non-fluorescent molecules in the compartment from which the flux flows.

$$R_I^f = \frac{N_c^f}{N_c^{tot}} R_I \quad (1c)$$

$$R_E^f = \frac{N_n^f}{N_n^{tot}} R_E \quad (1d)$$

Note that (1c) and (1d) hold independently of the kinetic equations for the original flux terms $R_I$ and $R_E$. Also assume that any spontaneous conversion between fluorescent and non- fluorescent molecules can be neglected. Then the differential equations for the fluorescent molecules are given by

$$\frac{d}{dt} N_c^f = -R_I^f + R_E^f = -\frac{N_c^f}{N_c^{tot}} R_I + \frac{N_n^f}{N_n^{tot}} R_E = -k_I N_c^f + k_{EV} N_n^f \qquad (1e)$$

$$\frac{d}{dt} N_n^f = +R_I^f - R_E^f = +\frac{N_c^f}{N_c^{tot}} R_I - \frac{N_n^f}{N_n^{tot}} R_E = +k_I N_c^f - k_{EV} N_n^f \qquad (1f)$$

where the first rewriting of the right hand-side of the ODEs made use of (1c) and (1d), and the second defined the two constants $k_i$ and $k_{EV}$. We name the second constant $k_{EV}$ and not $k_E$ since there is an inherent volume ratio between the compartments needed to convert concentrations into number of molecules (V was estimated experimentally in Figure 1 D). Note that eqs. (1e) and (1f) describe both the linear kinetics of equations 1a and b, and they provide a general formula for how the rate should be interpreted based on the original kinetics.

All in all, without making assumptions on the kinetics that regulates the import and export of the protein, the dynamics of the bleached and fluorescent fraction can be described by a simple two-compartment model using ordinary mass-action dynamics.

##### 1.2. The cytosol behaves as a homogeneous volume: comparison of shell models

One major assumption in our modeling approach is the homogeneity of the cytosol. That is, we assume that the proteins of interest are evenly distributed within this compartment, and that diffusion is fast enough that the probability of being transported into the nucleus is independent on the location in the cytosol. This is a valid approximation if the diffusion rate is much faster than the nuclear transport rates. If this were not the case, models that account for a more complex structure of the cytosol should give a better fit to the data. In this analysis we tested if models in which we subdivid the cytosol into several smaller compartments give significantly better fit than our simple two-compartment model.

We have approached this issue by constructing models where the interior of the cell is divided into a series of concentric shells. The nucleus corresponds to the innermost shell; in turn, it is surrounded by the innermost shell of the cytosol, which is surrounded by the next layer of the cytosol, and so on. In this analysis we have compared dividing the cytosol in 1 (the standard two-compartment model), 2 or 3 such shells. Each shell of the cytosol has been constructed to have equal volume (and therefore slightly different thickness).

The models have a diffusion constant, which in the models corresponds to the transport rate between neighboring shells (the same in both directions). This constant is higher between outer shells than inner shells, in proportion to the relative differences in surface area contacting the shells. The constant in the model is a free parameter, constrained to be larger than 1 and smaller than 10, and it's a multiplication factor of the first diffusion parameter. The optimization algorithm has chosen values between 1 and 8, depending on the data set. For the data set we show here, it was ~6.

We thus have the following models:

- 1-shell: 10 parameters (2 kinetic, 8 initial conditions, corresponding to the initial values of the fluorescent and dark species in each compartment after each photobleaching event)*
- 2-shell: 15 parameters (3 kinetic, 12 initial conditions)
- 3-shell: 20 parameters (4 kinetic, 16 initial conditions)

**In the main text we used a reduced version with 6 parameters, of which only four are initial conditions and correspond to the initial values of the fluorescent species only (Figure 1B).*

For each shell added, we add 1 diffusion parameter, and 4 initial-conditions parameters. This purposefully gives the model a lot of freedom, for example, we allow for it to completely change the state values between FRAP-trains in an unrealistic and discontinuous way. This is because if we can then reject these versions of the model, we can also reject versions where more care has been taken into choosing realistic initial conditions of the various shells.

We then contrasted these models to experimental data in the following way. We considered the FRAP experiments performed on YFP and Ace2 presented in the main paper, and classified our ~90 data sets into 5 different categories, based on the observed kinetics. These were: slow dynamics, medium dynamics, fast dynamics, linear response, and no response. From each category we chose two representative data sets, one that fitted well (i.e. had relatively low $\chi^2$cost), and one that fitted poorly (with relatively high $\chi^2$cost). We fitted the more complex models to the same data sets and observed the improvement in fit. We then tested if this improvement was significant using a standard likelihood ratio test. The results can be seen in the table below.

| Class | $\chi^2$Cost 1-shell | $\chi^2$ Decrease 2-shell | $\chi^2$ Decrease 3 -shell |
|---|---|---|---|
| Slow - high | 147.62 | -6.62 | -2.55 |
| Slow - low | 47.67 | -6.25 | -0.85 |
| Medium - low | 71.31 | -1.26 | -1.10 |
| Medium - high | 113.16 | -2.49 | -1.92 |
| Fast - low | 107.59 | -2.48 | -6.83 |
| Fast - high | 176.89 | -18.10 | -0.91 |
| Nothing - high | 317.26 | -8.13 | -0.53 |
| Nothing - low | 151.07 | -1.79 | -2.82 |
| Linear - low | 52.81 | -2.09 | -1.55 |
| Linear - high | 97.71 | -7.09 | -5.04 |
| *Cutoff* | | *11.07* | *11.07* |

Table S1. Shown are $\chi^2$ costs for the ten different data sets. First column shows the actual cost for the simplest model. The second and third columns show the successive decrease in $\chi^2$ cost obtained by adding a new shell. This value should be compared to the cutoff-value for a likelihood ratio test, which follows a $\chi^2$ distribution. Shown is the cutoff for 5% significance level. Significant results are highlighted in orange.

From our statistical analysis, we could only see a significant improvement in one of the cases (see Table S1). Note that with a 5% significance level we expect to see one false positive one time out of twenty, which is precisely the frequency we observed above. However, we chose to study the observed data set in more detail to see if we could understand how the more complex model achieved this better fit. The results are in Appendix Figure 1. The more complex model did not fit to the data in a qualitatively different way, rather it accomplished the improve cost by matching the initial conditions more flexibly.

In conclusion, one in twenty data sets showed an improvement in goodness of fit when going from a 1-shell model of the cytosol to a 2-shell model. However, for this data set, the improvement did not seem to come from a qualitatively different fit to the data. No data set showed any significant improvement when going from a 2-shell to a 3-shell model. Taken together, it does not seem that modeling the cytosol in more detail improves the overall goodness of fit.

### 2. Theoretical analysis on the impact of acquisition noise in the accuracy of the determined parameters

First, we simulated data similar to the one we obtained with our experimental protocol. To do this, we used the same ODEs 2-compartment model than for the fittings, with the time resolution of the experiments and "realistic" parameters and initial conditions (kI=0.4, kEV=4.41, $n_0$= 500, $c_0$=500). Then, we introduced noise according to the following equation

$$noisyData = simData + r * \frac{simData}{snr}$$

$$Data_{w/noise} = Data_{sim}\left(1 + \frac{r}{snr}\right) \tag{2a}$$

Where **snr** represents the signal to noise ratio (that was set to 1, 5, 10, 50 or 100) and **r** is a random number generated with the *randn* function of MATLAB, normally distributed with mean=0 and sd=1.

For each signal to noise ratio, we generated 10 datasets (Figure 1 C and Appendix Figure 2). When we studied the result of varying the number of FRAPs per train, we generated 10 independent datasets for each train length from 1 to 10 (Figure 1 C). Then, the simulated data was treated as experimental data.

### 3. FRAP images correction and quantification

Before fitting the model, we corrected the obtained experimental data for the non-cellular fluorescence background, cell's autofluorescence, imaging photobleaching (section 3.1, Appendix Figure 2), and reversion of the intentional photobleaching (section 3.2, Appendix Figure 2).

#### 3.1. Imaging photobleaching

In addition to the intended photobleaching of the FRAP experiments, some unintentional photobleaching also happened as a result of the acquisition of the images to follow the re-localization dynamics (i.e., imaging photobleaching). We minimized imaging photobleaching by opening the confocal pinhole as much as possible (500 µm, increasing photon counts at the expense of confocality) and reducing laser intensity to the minimum amount that resulted in an acceptable signal-to-noise ratio. Even under these conditions, imaging photobleaching was still detectable and we had to quantify it to correct the data (Appendix Figure 2 A). We performed this correction by introducing the experimentally determined photobleaching rate as a fixed parameter in the fitted model (see Section 4).

##### 3.2. Reversibility of photobleaching

Essential for a FRAP experiment is the assumption that any increase in fluorescence signal is due to movement of molecules to the bleached region and not from other processes, such as fluorophore maturation or reversibility of the photobleaching process. Maturation of YFP is too slow to have a significant effect during our FRAPs (half time for YFP maturation is ~ 40 min (Gordon et al., 2007), while the recovery is seen in 7 sec windows). However, it was important to determine and correct for the fraction, if any, of the photobleached YFP that could revert to the fluorescent state (Sinnecker et al., 2005; Henderson et al., 2007).

The reversibility of photobleaching was measured by completely photobleaching whole cells (so that fluorescence could not recover by molecules entering from other compartments) using laser intensities between 10 and 100% (Appendix Figure 2 B). For these experiments, cells were pre-treated with the translational inhibitor cycloheximide (CHX) for at least for 3 hours before the experiments, to allow all fluorescent molecules to mature (Gordon et al., 2007). We found that laser intensity did not change the characteristic time of recovery, but strongly changed the fraction of fluorescence recovered, following an exponential curve (Appendix Figure 2 B). Remarkably, for the YFP used here, 20% of the bleached molecules reverts to a fluorescent state after 1 min when photobleached with a 10% laser intensity, a number that reaches a minimum of 1% reversion with 50% laser intensity (Appendix Figure 2 B). In our experiments of trains of 4 FRAPs, the intentional photobleaching was performed with 100% laser intensity, so recovery can be neglected.

Similar experiments using cells fixed with 4% PFA showed no recovery (not shown). The discrepancy of these results can be explained if the fixating process also affects the recovery of the photobleaching and thus suggest that this control cannot be performed on fixed cells.

##### 3.3. Fluorescence quantification

To quantify total fluorescence in each compartment, we manually drew regions of interest (ROIs) in the nucleus and the cytosol and measured average fluorescence using the ImageJ plugin “Time Series Analyzer V2”. A typical ROI was a circle of 200 nm in radius (Appendix Figure 3 C). We studied the effect that drawing slightly different ROIs (different shapes, sizes and locations) would have on the estimated parameters, but we did not find any significant differences (not shown).

### 4. Fitting the nuclear transport model to the FRAP data

The data was fitted using an available R package, the core of which is the model presented in section 1. However, to better describe the experimental data, more parameters were introduced. These parameters were estimated independently from the nuclear transport rates, so they do not affect their determination. The complete correction and fitting procedure were able to adjust to the data in most cells, yielding well-determined kinetic rates for all the proteins studied (Figure 3 A).

#### 4.1. Modeling trains of FRAPs

From section 1, assuming steady-state before the first photobleaching event we obtain

$$\psi S_n^f(t) = \psi S_0^f \frac{\kappa_i}{\kappa_e+\kappa_i} \quad (4a)$$

$$\psi S_c^f(t) = \psi S_0^f \frac{\kappa_e}{\kappa_e+\kappa_i} \quad (4b)$$

$$\psi F_n^f(t) = \psi F_{n,0}^f \quad (4c)$$

$$\psi F_c^f(t) = \psi F_{c,0}^f \quad (4d)$$

for $t < t_1^{PB}$, where $t_1^{PB}$ is the time of the first photobleaching period, $\psi S_0^f$ the initial amount of total shuttling fluorescent protein (nuclear and cytoplasmic), $\psi F_{n,0}^f$ the initial amount of nuclear “fixed” fluorescent protein and $\psi F_{c,0}^f$ the initial amount of cytoplasmic fixed fluorescent protein ($\psi$ indicates that the variable is measured in units of fluorescence). Fixed protein refers to the molecules that for one reason or another are effectively immobile during the FRAP experiment (e.g., a fraction of the molecules might be bound to DNA in the nucleus).

Immediately after the i-th photobleaching event, the amount of shuttling proteins in each compartment will be given by

$$\psi S_{n,i}^f = (1 - PB_n)\, \psi S_n^f(t = t_i^{PB} - \Delta t) \quad (4e)$$

$$\psi S_{c,i}^f = (1 - PB_c)\, \psi S_c^f(t = t_i^{PB} - \Delta t) \quad (4f)$$

where $PB_n$ and $PB_c$ are the fraction of nuclear and cytoplasmic fluorescent proteins photobleached in each event, $t_i^{PB}$ the time of the i-th event and $\Delta t$ the sampling period. Note that $t = t_i^{PB} - \Delta t$ is the last time point before the photobleaching event. The dynamics of recovery for $t_i^{PB} \leq t < t_{i+1}^{PB}$ will be given by

$$\psi S_n^f(t) = \frac{\left(\psi S_{n,i}^f + \psi S_{c,i}^f\right)\ \kappa_i + \left(\psi S_{n,i}^f\, \kappa_e - \psi S_{c,i}^f\, \kappa_i\right) e^{-(\kappa_e+\kappa_i)\,t}}{\kappa_e+\kappa_i} \quad (4g)$$

$$\psi S_c^f(t) = \psi S_{n,i}^f + \psi S_{c,i}^f - \psi S_n^f(t) \quad (4h)$$

For the fixed fraction, the nuclear and cytoplasmic amounts after the i-th photobleaching event will be given by

$$\psi F_n^f(t) = (1 - PB_n)^i\, \psi F_{n,0}^f \quad (4i)$$

$$\psi F_c^f(t) = (1 - PB_c)^i\, \psi F_{c,0}^f \quad (4j)$$

##### 4.2. Parameterization of focus drift

The functions $\phi_n(t)$ and $\phi_c(t)$ describe the smooth changes in nuclear and cytoplasmic focus due to focus drift. If the focus remained constant during all the experiment then $\phi_n(t) = \phi_c(t) = 1\ \forall\ t$, but the focus can change slightly during the experiment, which produces changes in the observed fluorescence intensity. This effect could be neglected for the relatively large cytoplasm (in comparison to the slicing of the confocal microscope), but it produced measurable differences for the nucleus and therefore has to be accounted for in the model. Therefore, in what follows we considered $\phi_c(t) = 1\ \forall\ t$. To avoid identifiability issues between the FRAP and focus-drift dynamics, we constrained $\phi_n(t)$ to remain constant during a FRAP and to vary smoothly between consecutive FRAPs.

$$\phi_n(t) = \phi_{n,j}\ \ if\ \ t_j \le t < t_{j+1} \quad (4k)$$

$$\phi_{n,j} = 1 + \alpha_{n,1}(t_j - t_0) + \alpha_{n,2}(t_j - t_0)^2 \quad (4l)$$

where j denotes the FRAP in the series, $t_j$ the start time of each FRAP and the parameters $\alpha_{n,1}$ and $\alpha_{n,2}$ are fitted.

##### 4.3. Geometric volume fraction and autofluorescence

Geometric volumes were experimentally determined for each cell by fitting ellipses to the nucleus, vacuole and the whole cell (Appendix Figure 3 B), and calculating the corresponding volumes ($\tilde{V}_n^{geom}$, $\tilde{V}_{vacuole}^{geom}$ and $\tilde{V}_{cell}^{geom}$), assuming height equal to the minor axes. Uncertainties in volume estimations ($\Delta\tilde{V}_n^{geom}$ and $\Delta\tilde{V}_c^{geom}$) were in the order of 10% as assessed by multiple independent estimations using this method. The cytoplasmic geometric volume was estimated as the cell's volume minus the volumes of the nucleus and vacuole $\left(\tilde{V}_c^{geom} = \tilde{V}_{cell}^{geom} - \tilde{V}_n^{geom} - \tilde{V}_{vacuole}^{geom}\right)$. The estimated value for the geometric volume ratio $\tilde{V}_{frac}^{geom} = \frac{\tilde{V}_c^{geom}}{\tilde{V}_n^{geom}}$ was calculated and used to constrain the parameter $V_{frac}^{geom}$ during the fitting procedure.

We considered equal autofluroescence levels between compartments $\psi^{auto} \equiv \psi_c^{auto} = \psi_n^{auto}$, as we could detect no differences between compartments in the parental (unlabeled) strains.

##### 4.4. Parameter estimation and optimization

The following table describes all the parameters in the model.

| Parameter | Description | Initial value |
|---|---|---|
| $\boldsymbol{log(\kappa_i)}$ | Partially adimensional nuclear import rate $\kappa_i = \frac{k_i}{V_c^{eff}}$ | logarithmic grid |
| $\boldsymbol{log(\kappa_e)}$ | Partially adimensional nuclear export rate $\kappa_e = \frac{k_e}{V_n^{eff}}$ | logarithmic grid |
| $\boldsymbol{\psi S_{tot,0}^f}$ | Total initial shuttling fluorescent protein | Estimated from time series |
| $\boldsymbol{\psi F_{n,0}^f}$ | Initial nuclear fixed fluorescent protein | Estimated from time series |
| $\boldsymbol{\psi F_{c,0}^f}$ | Initial cytoplasmic fixed fluorescent protein | 0 |
| $\boldsymbol{PB_n}$ | Fraction of nuclear proteins photobleached | Estimated from time series |
| $\boldsymbol{PB_c}$ | Fraction of cytoplasmic proteins photobleached | Estimated from time series |
| $\boldsymbol{\psi^{auto}}$ | Autofluorescence level | $0.1\ \psi S_{tot,0}^f$ |
| $\boldsymbol{\alpha_{n,1}}$ | Nuclear focus-drift linear coefficient | 0 |
| $\boldsymbol{\alpha_{n,2}}$ | Nuclear focus-drift quadratic coefficient | 0 |
| $\boldsymbol{log(V_{frac}^{geom})}$ | Geometric volume fraction | Estimated from image |

Table S2: Parameters and initialization values of the full ODE model used to fit the train of FRAPs

Initial values for the partially adimensional transport rates $\kappa_i$ and $\kappa_e$ were scanned logarithmically from 0.01 to 10 $s^{-1}$ . All other initial values of the parameters were either fixed or estimated from the images. Optimization was done using nonlinear least-square solver (lsqnonlin). The cost function minimized was defined as

$$Cost(\vec{p}) = \sum_{k=1}^{T} \frac{\left(\psi_{n,\vec{p}}(t_k) - \tilde{\psi}_n(t_k)\right)^2}{\sigma_n^2} + \sum_{k=1}^{T} \frac{\left(\psi_{c,\vec{p}}(t_k) - \tilde{\psi}_c(t_k)\right)^2}{\sigma_c^2} + \frac{\left(log(V_{frac}^{geom}) - log(\tilde{V}_{frac}^{geom})\right)^2}{\left(\Delta log\left(\tilde{V}_{frac}^{geom}\right)\right)^2} \qquad (4m)$$

where the sums are over the $T$ time frames fitted (spanning the train of FRAPs), $\vec{p}$ is the parameter vector (see Table S2), and $\tilde{\psi}_n$ [$\tilde{\psi}_c$] denote the experimentally measured values for nuclear [cytoplasmic] fluorescence (corrected for imaging photobleaching). The standard deviations for each compartment, $\sigma_n$ and $\sigma_c$, were calculated per cell from movies in which the pertinent cell was not being photobleached (when its mother or daughter was being photobleached) (Appendix Figure 3 B). To this end, we detrended the data using a loess model to remove putative focus drift or imaging photobleaching effects. The last term in the cost function penalizes for deviations of the parameter $V_{frac}^{geom}$ from the experimentally estimated value $\tilde{V}_{frac}^{geom}$ in logarithmic scale. The denominator of this term evaluates to the square of the relative error of $\tilde{V}_{frac}^{geom}$.

$$\left(\Delta log\left(\tilde{V}_{frac}^{geom}\right)\right)^2 = \left(\frac{\Delta\tilde{V}_{frac}^{geom}}{\tilde{V}_{frac}^{geom}}\right)^2 = \left(\frac{\Delta\tilde{V}_{n}^{geom}}{\tilde{V}_{n}^{geom}}\right)^2 + \left(\frac{\Delta\tilde{V}_{c}^{geom}}{\tilde{V}_{c}^{geom}}\right)^2 \qquad (4n)$$

##### 4.5. Nuclear fixed fraction pre-estimation

In order to prevent identifiability problems for $k_I$ and $k_E$ resulting from the inclusion in the model of the free parameters for the initial conditions of the nuclear and cytoplasmic fixed fractions, we pre-estimated the initial value of these fractions running a different analysis on the experimental data. The general idea for this analysis stems from our previous work {Doncic et al., 2015} and is based on the notion that following the recovery after photobleaching of a compartment, the mobile and the fixed fractions behave differently: while the mobile nuclear fraction of a fluorescent protein reaches the same relation with the other compartment (Sn/Sc), the fixed nuclear fraction (fn/fc) is irreversibly reduced (within that time window). To calculate the fraction of shuttling molecules and the amount of fixed fluorescence, this analysis then compares the initial (right before photobleaching) and final (in equilibrium) nuclear-to-cytoplasmic fluorescence ratio. If they are not equal, it implies that the fraction of fixed molecules is non-zero. Note that the analysis just evaluates these two steady-state time points and not the entire dynamics of the time-series. In this way, it is possible to estimate the number of fixed molecules in the bleached compartment.

Appendix Figure 4 A shows the results of simulations implemented on COPASI to illustrate the above concept. This simple model considers 2 compartments -nucleus and cytoplasm-, and 1 species, that can shuttle in and out, or be fixed in the nucleus. It assumes that the rates for the fixating reactions happen in a much slower time scale than nuclear shuttling and that there is no imaging photobleaching. Trains of FRAPs were modeled as a fractional reduction in the number of nuclear molecules, no matter if they were fixed or mobile. The model clearly shows how the presence of a nuclear fixed fraction reduces the nucleus/cytoplasm ratio after photobleaching (Appendix Figure 4 A).

Following this idea, we derived an expression for the estimation of the nuclear fixed molecules from the initial and final ratios before performing the fitting of the time-series. The equivalence of the initial and final nuclear-to-cytoplasmic fluorescence ratio of total fluorescence can be expressed as:

$$\frac{T_n^f(t=i)}{T_c^f(t=i)} * r = \frac{T_n^f(t=f)}{T_c^f(t=f)} \qquad (4o)$$

Where r is the fraction of the total fluorescence reduced in the photobleaching, $T^f$ represents the total amount of fluorescent molecules, the sub-indexes n and c denote the nuclear or cytosolic compartments, and I and f indicate the initial and final times, respectively. Note that all these $T^f$ can be easily measured from the initial and final images, and thus r can be easily estimated as well.

Then, the $T^f$ can be broken into its mobile (S) and the fixed (F) components.

$$\frac{F_n^f(t=i)+S_n^f(t=i)}{F_c^f(t=i)+S_c^f(t=i)} * r = \frac{F_n^f(t=f)+S_n^f(t=f)}{F_c^f(t=f)+S_c^f(t=f)} \qquad (4p)$$

If we assume that there are no fixed molecules in the cytoplasm, this becomes:

$$\frac{F_n^f(t=i)+S_n^f(t=i)}{S_c^f(t=i)} * r = \frac{F_n^f(t=f)+S_n^f(t=f)}{S_c^f(t=f)} \qquad (4q)$$

And, assuming that after the 4 FRAPs the nuclear fixed fraction would be completely photobleached

$$\frac{F_n^f(t=i)+S_n^f(t=i)}{S_c^f(t=i)} * r = \frac{S_n^f(t=f)}{S_c^f(t=f)} \qquad (4r)$$

If at the final time point the mobile fraction has reached steady-state $\frac{S_n^f(t=i)}{S_c^f(t=i)} = \frac{S_n^f(t=f)}{S_c^f(t=f)}$, and replacing in 5q

$$F_n^f(t=i) = \frac{S_n^f(t=i)}{r} - S_n^f(t=i) \qquad (4s)$$

Expressing equation 4r as a function of the total fluorescence (Tn) form eq 4o, (what may be measured experimentally),

$$F_n^f(t=i) = T_n^f(t=i) - T_n^f(t=i) * r \qquad (4t)$$

We get the expression that was used for the estimation of the initial condition. During the fitting this value was optimized with boundaries that accepted a variation of ±20%. As a result of the pre-estimation of the nuclear fixed molecules, the adjustment of the model to the data improved, resulting in optimized parameters with smaller uncertainties (Appendix Figure 4 B).

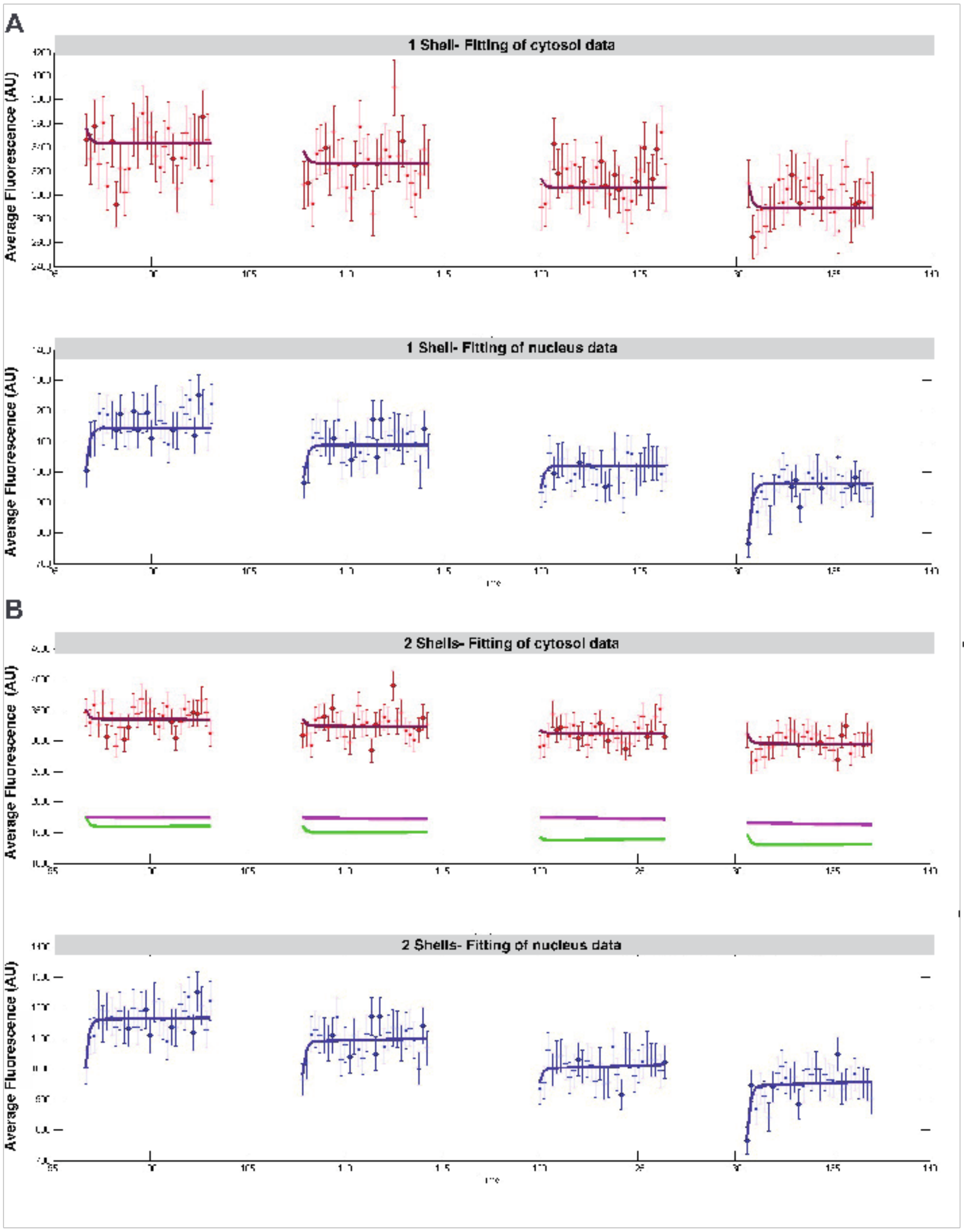
A
1 Shell- Fitting of cytosol data
Average Fluorescence (AU)
1 Shell- Fitting of nucleus data
Average Fluorescence (AU)
B
2 Shells- Fitting of cytosol data
Average Fluorescence (AU)
2 Shells- Fitting of nucleus data
Average Fluorescence (AU)

**Appendix Figure 1. Comparison between a 1-shell (A) and 2-shell** (B) model of the cytosol for the data set that showed an improved goodness of fit. Red error bars show the observed cytosolic data, blue error bars show the observed data in the nucleus. Dark red and dark blue continuous lines show model fits to this data for the 1-shell and 2-shell model top and bottom respectively. Continuous green and magenta lines in bottom figure show the time course for the inner and outer shell respectively in the 2-shell model.

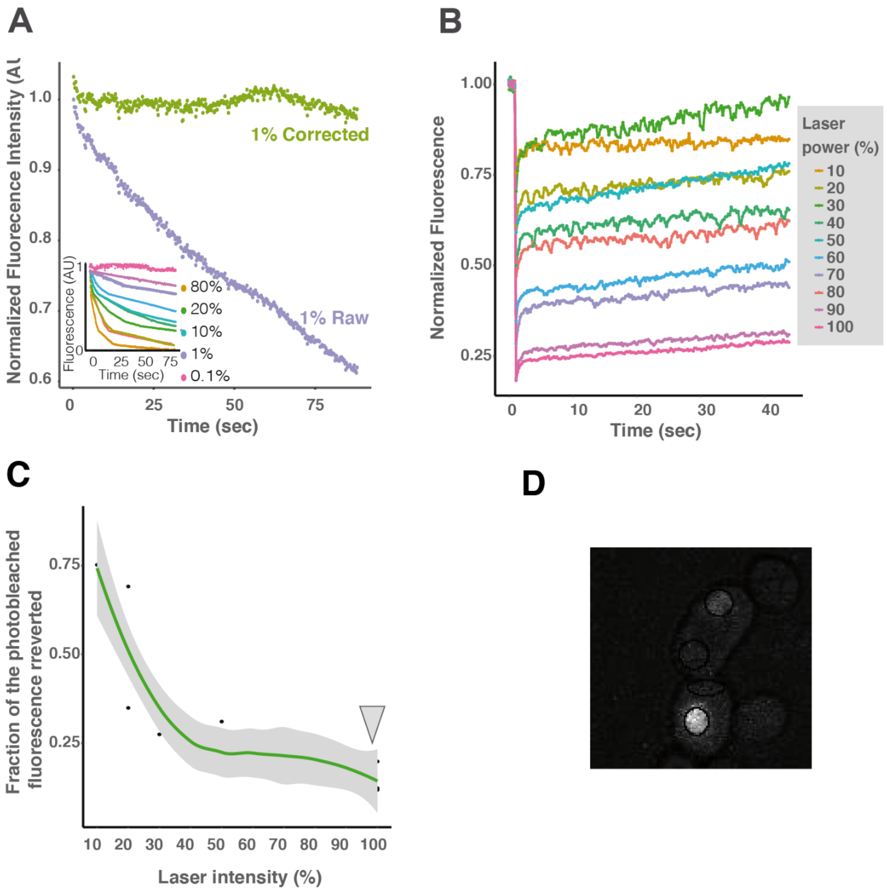


**Appendix Figure 2. Imaging conditions characterization for the FRAP experiments.** (A) Correction of imaging photobleaching. After 3 hs cycloheximide (CHX) treatment, 400 consecutive images were taken, under the same conditions than those used for FRAP experiments (purple dots). This data was used to implement a correction for imaging photobleaching in the FRAP experiments (the result is shown in green dots). Inset: imaging photobleaching for different laser intensities. The average of two measurements is shown for each. (B) Characterization of the reversion of photobleached YFP fluorescence when excited with a 515 nm laser. Yeast expressing YFP were treated with CHX and analyzed 3 hs later to ensure full fluorophore maturation. The entire yeast was photobleached with the indicated laser intensity. The data was fitted by an exponential curve with parameters: lambda=54,036, A=0,672 y B= 0,0548. (C) The recovered fraction (the fraction was computed as (final fluorescence - fluorescence after photobleaching)/ (initial  fluorescence- fluorescence after photobleaching), so it represents the fluorescence recovered/ fluorescence lost) as a function of laser intensity is shown. The gray shading depicts the 95% confidence interval from 3 repetitions per laser power. YFP photobleaching is irreversible when performed with high laser power. (D) Example of a mother-daughter yeast pair of cells with the ROIs measured. Fluorescence corresponds to YFP-Ace2.

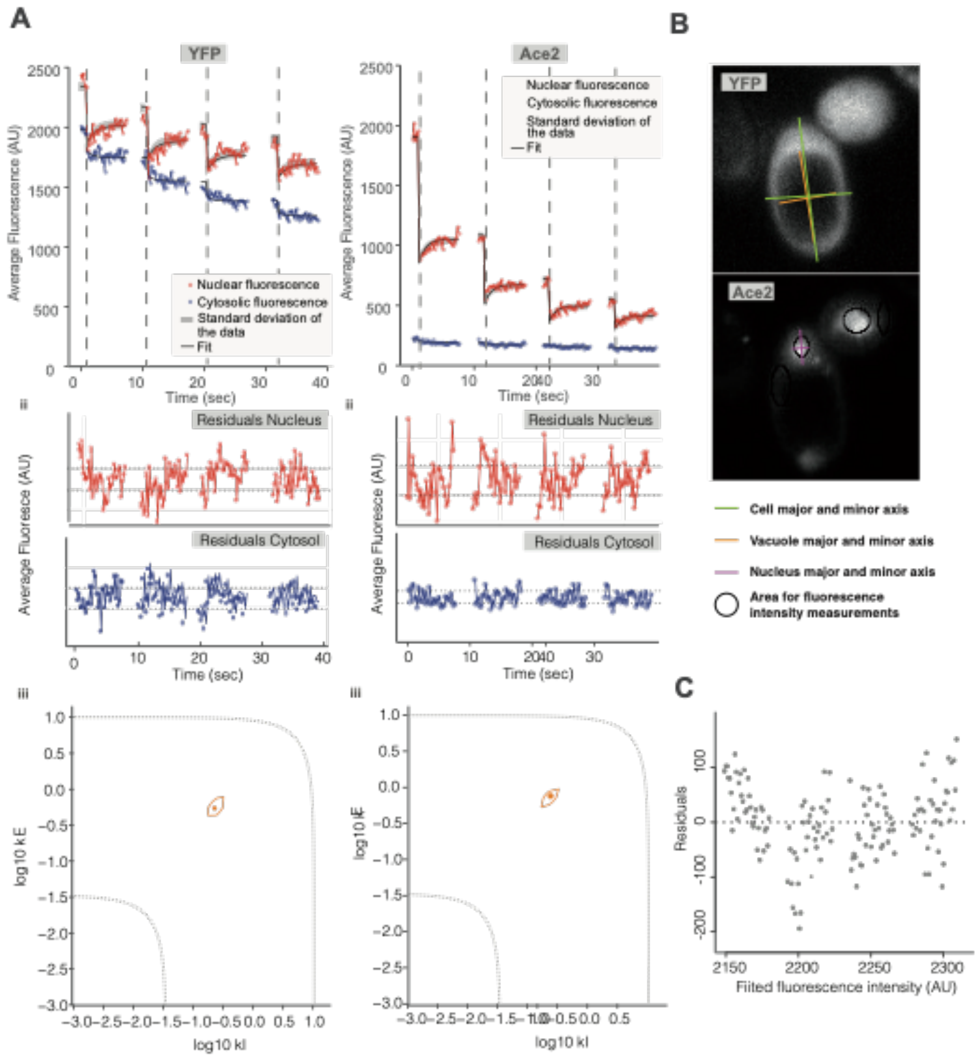
A
B
C
YFP
Ace2
Nuclear fluorescence
Cytosolic fluorescence
Standard deviation of the data
Fit
Average Fluorescence (AU)
Time (sec)
ii
Residuals Nucleus
Residuals Cytosol
Average Fluoresce (AU)
iii
log10 kE
log10 kI
Cell major and minor axis
Vacuole major and minor axis
Nucleus major and minor axis
Area for fluorescence intensity measurements
Residuals
Fitted fluorescence intensity (AU)

F

**Appendix Figure 3. Experimental determination of nuclear transport rates in single cells.** (A) Examples of fitted data. Left: free YFP, right: Ace2-YFP. i: Fluorescence intensity curves (gray shading depicts the standard deviation of the experimental data) and fittings (black lines). ii: Residuals of the adjustments. iii: Estimated import and export rates. The full line limits all acceptable values ($p<0.05$). Both nuclear transport rates can be identified, and thus estimated independently. The dotted lines show the limits of the parameter values that can be estimated with confidence using our experimental protocol (time resolution, duration of the experiment). (B) Example of the geometric measurements of the cells that were used as an input for the optimization procedure. (C) Estimation of the standard deviation "sigma" of the signal under the FRAP imaging conditions

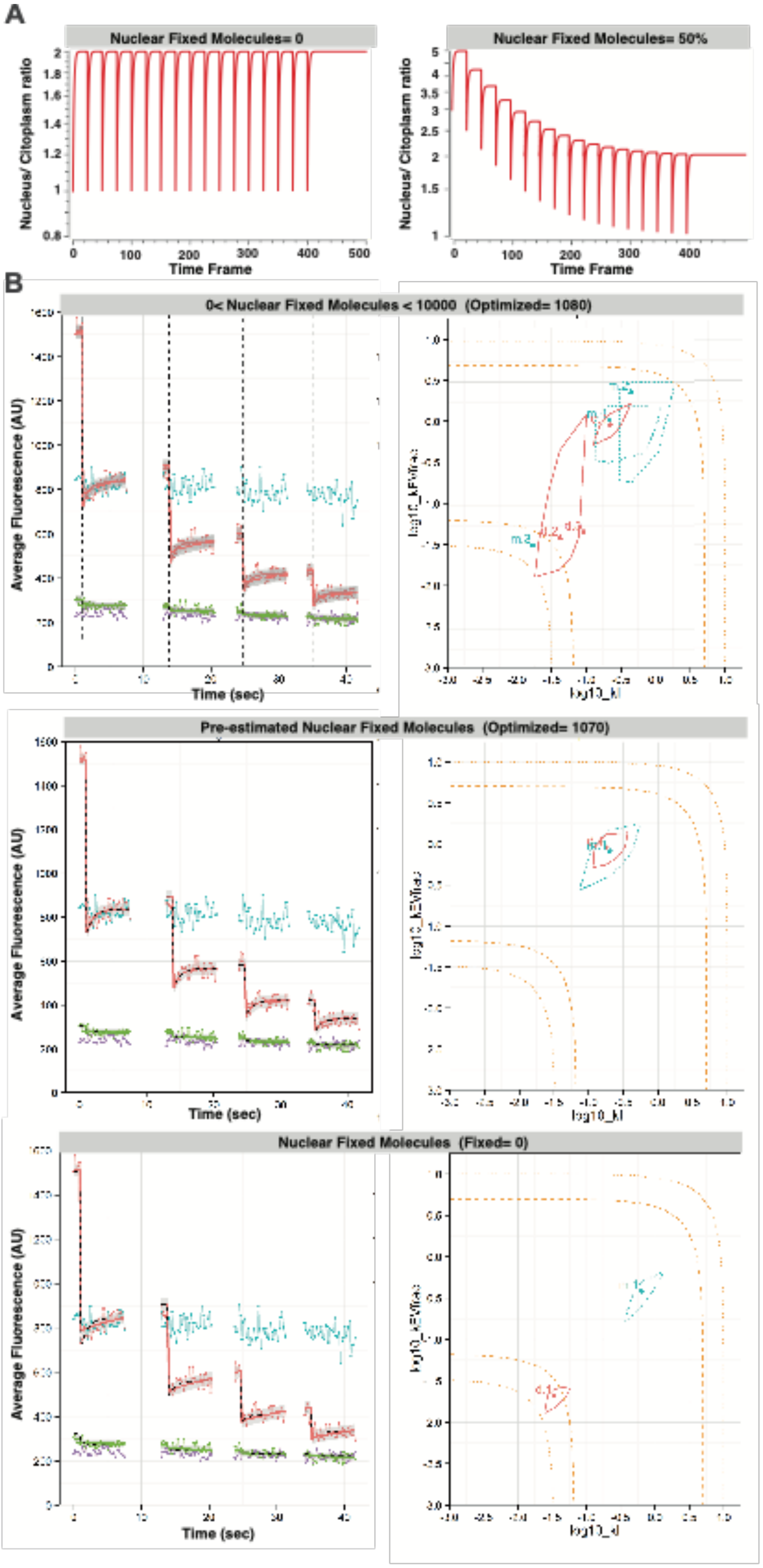
A
Nuclear Fixed Molecules= 0
Nuclear Fixed Molecules= 50%
Nucleus/ Citoplasm ratio
Time Frame
B
0< Nuclear Fixed Molecules < 10000 (Optimized= 1080)
Average Fluorescence (AU)
Time (sec)
log10_kEWfree
log10_kl
Pre-estimated Nuclear Fixed Molecules (Optimized= 1070)
Nuclear Fixed Molecules (Fixed= 0)

**Appendix Figure 4. Estimation of the nuclear fixed fraction from steady-state images pre and post photobleaching of the nucleus.** (A) Simulations where a train of 17 partial FRAPs were performed on cells that either had no fixed molecules in the nucleus or had 50% of the total molecules fixed in the nucleus. No cytoplasmic fixed molecules were introduced. When a fixed fraction of molecules is included, the steady-state nucleus/cytoplasm ratio is reduced with increasing FRAPs. (B) Example of fitting of an Ace2 FRAP experiment, letting the nuclear fixed molecules parameter (Fn0) free (top), having it pre-estimated according to section 4 of the Appendix (middle), or considering no fixed molecules in the nucleus (bottom). The plots are as in Appendix Figure 3. In the right panels, the pink lines show acceptable fitting for the daughter cell and the light-blue show the acceptable values for the mother cell. Note that in the top example, problems with the determination of the kinetic nuclear transport parameters arise. These issues are seen as larger areas of acceptable parameters (right), and the presence of several minimums. When no fixed fraction was introduced, the model does not adjust the data well (bottom, left).
